# Disease progression shapes circadian and locomotor disruption in a model of multiple sclerosis

**DOI:** 10.64898/2026.10.08.757675

**Authors:** Vina W. Li, Julia P. Segal, Nader Ghasemlou

**Author notes:** Corresponding Author: Nader Ghasemlou, PhD Krembil Brain Institute, University Health Network, 60 Leonard Ave, suite 7KD-418 Toronto, Ontario, Canada M5T 0S8.

## Abstract

Multiple sclerosis (MS) is an autoimmune demyelinating disease of the central nervous system featuring a wide range of debilitating symptoms. Wearable activity recording devices have recently been adopted to improve symptom monitoring in MS, capturing symptoms related to sleep/wake cycles, circadian disruption, and fatigue. However, comparable measures of activity have yet been established in pre-clinical models of MS, the most widely used being experimental autoimmune encephalomyelitis (EAE). We therefore sought to characterize changes in continual home-cage activity over the course of EAE, and establish whether home-cage monitoring can be used as an observer-independent measure of disease progression. Our results showed that locomotor activity in the mid-active/dark phase was most severely affected by EAE. Disrupted circadian (24-hour) activity was observed during the peak and onset stages of disease. Bout activity analysis revealed >40-minute long activity bouts as a sensitive indicator of non-motor symptoms independent of clinical score-based motor deficits. Current assessment of disease progression in EAE relies on clinical scores, which primarily assess motor deficits through visual observation prone to observer bias. Various computed activity parameters were found to be altered during key disease stages, suggesting that a combination of activity measures can serve as objective indicators of disease progression. Overall, our results show that home-cage activity effectively captures signs of non-motor symptoms in the EAE model of MS and can serve as an observer-independent measure of EAE disease progression.

## 1. INTRODUCTION

Multiple sclerosis (MS) is an autoimmune disease of the central nervous system (CNS) characterized by inflammatory myelin lesions and chronic neurodegeneration. People living with MS (pwMS) experience a wide range of debilitating symptoms, including altered sleep/wake cycles and fatigue (Crayton et al., 2004). Over 50% of pwMS frequently experience insomnia, daytime somnolence and nighttime awakenings (Bamer et al., 2008, Braley and Boudreau, 2016, Kouchaki et al., 2025, Sakkas et al., 2019). Exhaustion and low motivation are fatigue-related symptoms that often co-occurr with disrupted sleep/wake cycles (Attarian et al., 2004, Bhattarai et al., 2023, Taphoorn et al., 1993) and reduced physical activity (Gashi et al., 2024, Aboagye et al., 2025). Sleep dysregulation (Fang et al., 2019, Nutt et al., 2008) and low physical activity (Schuch et al., 2017, Stubbs et al., 2016) are also common among pwMS (Feinstein et al., 2014, Pucak et al., 2007). An important aspect of MS pathology is inflammation in both central and peripheral nervous systems (Haase and Linker, 2021). Inflammatory cytokines such as IL-1β and TNF-α can impair normal functions of the hypothalamus-pituitary-adrenal (HPA) axis (Dunn, 2000, Turnbull and Rivier, 1995), pineal gland (Carvalho-Sousa et al., 2011, Markus et al., 2018) and reward circuits (Felger and Treadway, 2017, Vichaya and Dantzer, 2018), disrupting mechanisms affecting arousal, sleep and motivation.

Symptoms of MS also feature high levels of variability both between and within individuals (De Somma et al., 2018). Beyond well-documented differences in disease progression (Engelhardt et al., 2022), studies using ecological momentary assessment have shown daily and often moment-to-moment fluctuations in MS symptom severity (Powell et al., 2017, Kratz et al., 2017a, Kratz et al., 2017b). Current symptom assessment in MS involve standardized rating scales, self-report questionnaires and performance testing during clinical visits (Cutter et al., 1999, Kos et al., 2005, Kurtzke, 1983, Miller et al., 2000), leaving symptom fluctuations uncaptured. These limitations have driven a shift towards continuous activity monitoring using wearable sensors, which capture symptom fluctuations in real-life settings that could complement current clinical assessments (Adao Martins et al., 2021, Alexander et al., 2021). However, no comparable methods of activity tracking currently exist in pre-clinical models of MS.

Experimental autoimmune encephalomyelitis (EAE) is a widely used animal model of MS wherein mice experience inflammatory demyelination and ascending paralysis similar to relapse-remitting MS (Constantinescu et al., 2011). The EAE model also recapitulates the disease variability seen in MS, with documented between-animal variations in symptom severity, disease progression and incidence (Simmons et al., 2013, Lee et al., 2003, Hasselmann et al., 2017, Berard et al., 2010, Mayrhofer et al., 2021). The majority of pre-clinical studies focus on neuroinflammation, demyelination and motor deficits, with limited investigation of the behavioural impact of EAE pathology. The impact of EAE progression on function and behaviour is reflected through locomotor activity count (Grace et al., 2017, Jones et al., 2008, Rahn et al., 2014), grip strength (Jones et al., 2008, Rahn et al., 2014, Osorio-Querejeta et al., 2022, Ondek et al., 2021), anxiety behaviour (Faraji et al., 2022) and thermoregulation (Faraji et al., 2022). However, these assays provide only a snapshot at the time of measurement, and are often carried out in unfamiliar testing environments that can induce stress-related changes (Gouveia and Hurst, 2017, Sturman et al., 2018). Although there is evidence of altered HPA axis activity (Trifunovic et al., 2021) and clock gene expression (Buenafe, 2012) in the EAE model, sleep/wake behaviour and circadian activity have yet been assessed.

There is a pressing need for observer-independent measures of disease progression that also capture longitudinal spontaneous activity in EAE model, with an over-reliance on clinical scoring systems to assess changes in functional outcomes that primarily evaluate motor deficits by visual observation (Palle et al., 2016). These scales are prone to inter-rater variability (Palle et al., 2016); compounded with the inherent variability of the model, reliance on subjective scoring scales could compromise the reproducibility of EAE studies and leave non-motor but clinically relevant symptoms undetected. Recently, home-cage monitoring has emerged as a valuable tool for assessing laboratory rodent activity across disease models (de Visser et al., 2006, Fisher et al., 2012). The longitudinal data collected can be used to detect changes in spontaneous activity over extended periods of time, ideal for characterizing behavioural outcomes through the course of disease. Here, we sought to determine whether home-cage activity data can be used as a monitoring tool for the progression of EAE and detection of continuous changes in spontaneous activity. Our work suggests that this tool provides a comprehensive means of capturing both motor and non-motor outcomes in EAE, with a more nuanced analysis of functional outcomes than standard clinical scoring scales. Thus, continuous monitoring of home-cage activity, which is increasingly adopted by animal facilities, may provide objective, reproducible, and quantifiable measures of disease to facilitate research in pathological mechanisms and therapeutic development.

## 2. METHODS

### 2.1 Animals

Female C57BL6/J mice (JAX #000664) were bred in our facility, with temperature controlled at 23°C and lights on a 12:12 light/dark cycle (lights on/off at 07:00 and 19:00); light changes were instantaneous and without ramped transitions. Access to standard mouse chow and water was provided *ad libitum*. Mice were housed n=2 per cage in the Digital Ventilated Cage (DVC) system (DGM500; Tecniplast, Italy) that registers fluctuations in electromagnetic fields caused by animal movements (Klein et al., 2022)for home-cage activity recording in a standard housing rack and cage system. Animal husbandry and supportive care were performed within 1-hour after lights-on or 1-hour before lights-off to minimize disturbance to normal circadian and locomotor activity. The use and care of animals in this study was completed in accordance with the Canadian Council on Animal Care and ARRIVE guidelines under Queen’s University Animal Use Protocol #2428.

### 2.2 EAE induction and scoring

Mice were lightly anesthetized with 1-2% isoflurane at 8-10 weeks of age to induce EAE and for sham controls. All mice received intraperitoneal injections of pertussis toxin (PTX; 75ng, Hooke Laboratories, USA; or 200ng, List Laboratories, USA) at time of immunization and again 48 hours later. Pertussis toxins from different suppliers and lots were titrated to match EAE disease severity. Immunization was performed through hind flank subcutaneous injections with a 1:1 emulsion of myelin oligodendrocyte glycoprotein (MOG; 50ug/ml, Synpeptide, China) and complete Freud’s adjuvant (CFA; Millipore Sigma, USA; 4mg/ml). Sham mice were injected with an emulsion of phosphate buffered saline and CFA, without MOG. Clinical score (CS) was used to assess disease progression daily using a 5-point clinical score scale (0 – no symptoms, 1 – limp tail, 2 – hind limb weakness, 3 – hind limb paralysis, 4 – front limb paralysis, 5 – moribund; half points were assigned between grades) (Miller and Karpus, 2007). After onset of disease, mice were provided with supportive care including moist chow on cage floor, saline injection and bladder expression where needed. Green clay was used to treat injection site lesions, where necessary.

### 2.3 Home cage activity data collection and analysis

The Tecniplast Digital Ventilated Cage (DVC) system registers fluctuations in electromagnetic fields caused by animal movements (Klein et al., 2022) and was used to collect animal home-cage activity data. The system uses an electromagnetic baseplate installed below each individual cage, with each baseplate having 12 electrodes that generate a 12-zone electromagnetic field on the cage floor. The capacitance of electromagnetic fields is altered when movements of water-rich objects (i.e. mice) are present in the field (Klein et al., 2022). Depending on the amount of movement, the activity data is stored as an activity index per minute and is accessible through the DVC Analytics (Tecniplast, Italy) platform. Mice were placed into DVC cages two weeks before immunization and recordings ending on 28 days post-immunization (dpi). Average activity from the 5 days prior to immunization was used as baseline activity (shown as “BL”) and average activity of 0-2 dpi was used as injection day activity (shown as “INJ”). Cage data from different sets of mice were matched by extracting data from 5 days pre-immunization to 27dpi, or the last day with complete 24-hour data before endpoint. Data analysis in this study is conducted using R v4.5.1 and ClockLab Analysis (Version 6; Actimetrics, USA). Packages used in R include dplyr v1.1.4 (Wickham, 2023) and ggplot2 v3.5.2 (Wickham, 2016).

### 2.4 Locomotor activity data processing

Activity index analysis was performed using indices assigned by the DVC system on movements detected per minute. Time is denoted using zeitgeber time (ZT), where ZT0 is lights-on time (07:00) and ZT12 is lights-off time (19:00). Activity index per minute was used to generate heatmaps of activity. For locomotor activity analysis per disease stage, the average activity of all days in each disease phase was used. To investigate activity distribution over 24 hours in each disease phase, a 60-minute average smoothing was performed on the average activity of disease stage to reveal trends that would otherwise be obscured by random activity fluctuation.

### 2.5 Bout activity analysis

Activity indices were transformed into binary data frames. If activity > 0, the minute is considered active and assigned a value of “1”. If activity = 0, the minute is considered inactive and assigned a value of “0”. Consecutive active minutes separated by at least 1 inactive minutes were considered an activity bout. Activity bouts were sorted based on their lengths into 6 bins (1-minute, 2-5 minutes, 6-10 minutes, 11-20 minutes, 21-40 minutes and >40 minutes). *Activity bout duration* was calculated as the total number of active minutes in each bin. *Activity bout frequency* was calculated as the number of activity bouts in each bin. *Activity bout index* was calculated as the sum of activity index of all activity bouts in each bin. *Normalized activity bout duration* was calculated by dividing the total activity bout duration in a bin by the total activity bout duration of all bins. *Normalized activity bout frequency* was calculated by dividing the number of activity bouts in a bin by the total activity bout frequency of all bins. *Normalized activity bout index* was calculated by dividing the sum of activity bout index in a bin by the total activity index of all bins. Rest bout parameters, including *rest bout duration, rest bout frequency*, *normalized rest bout duration* and *normalized rest bout frequency*, were calculated in the same manner except for counting the inactive minutes instead. For disease stage analysis, bout measures were generated from the total duration of each disease stage. For daily bout analysis, bout activity was computed from every consecutive 24 hours of activity. The cut-off time for every 24 hours of activity was set at ZT3, or after the ZT21-2^+1^ high activity period. As long activity bouts predominantly take place in the high activity periods, cut-off time at a low activity period prevents capturing artifacts caused by interrupting long activity bouts.

### 2.6 Circadian rhythm analysis

Activity data was converted to .AWD format to allow for analysis with ClockLabs software. Non-parametric parameters were used to analyze circadian rhythms of locomotor activity. *Relative amplitude* measures the distinction between active and resting phases using activity from the most active 10 hours and least active 5 hours. *Intradaily variability* measures the integrity of rest and active phases, or activity fragmentation. *Periodicity* was assessed using the Lomb-Scargle periodogram (Refinetti et al., 2007). The normalized power of the near 24-hour peak was used as a measure of changes in periodicity with disease, instead of an indication of the actual activity period. Where the peak near 24-hour is below significance (p > 0.05), normalized power of the dominant peak between 23.5-hour and 24.5-hour was used (Brown et al., 2019, Blume et al., 2017). Activity of every consecutive 24-hour period was used to generate daily circadian rhythm measurements. The overall activity duration of each disease stage was used to generated disease stage-specific circadian measurements.

### 2.7 Statistical analysis

Statistical comparison was performed using Prism 10 (GraphPad, California, USA). For clinical score and activity comparison between EAE and sham mice, two-way repeated-measure ANOVA was used followed by post-hoc Bonferroni’s analysis. For clinical score and activity comparison among EAE, atypical EAE and sham mice, two-way repeated-measure ANOVA was used followed by post-hoc Tukey’s analysis. For comparison between bout activity frequency in different disease stages in EAE mice, paired Student’s t-test was used. For comparison of bout activity in each disease stage among EAE, atypical EAE and sham mice, one-way ANOVA with post-hoc Tukey’s analysis was used. Statistical comparison details are included in Supplementary Table 1.

## 3. RESULTS

### 3.1 Mice exhibit time-of-day dependent reduction in locomotor activity over the course of EAE

Home-cage activity was recorded for 5 days after a one-week habituation to establish baseline measurements prior to EAE induction. Activity was monitored throughout the disease course and clinical score was used to assess daily change in motor deficits (Fig. 1A). The average clinical score profile of all EAE animals was used to determine disease stages for activity analysis. The pre-onset phase of disease was defined as from the day mice have recovered from disease induction and to the appearance of motor symptoms (i.e. 5-9dpi; average clinical score: 0.00). The onset phase was characterized by a period of consistent increase in clinical scores (i.e. 10-14dpi; average clinical score: 0.91), which then plateaued during the peak phase of disease (i.e. 15-20dpi; average clinical score: 2.35). The simultaneous reduction and plateau of clinical scores marked the chronic phase of disease (i.e. 21-27dpi; average clinical score: 1.87) (Fig. 1B; two-way RM ANOVA; interaction [CS x dpi]: F(1.815, 36.30) = 53.50, p < 0.0001). To characterize changes in home-cage activity, we performed locomotor and rest/active phase structural analysis. A heatmap of activity index showed a general reduction in the activity of EAE mice compared to sham controls after disease onset, with a partial recovery in the chronic phase (Fig. 1C). Change in home-cage activity was consistent with the progression and recovery of motor deficits in EAE, with activity levels inversely correlated with motor deficit severity. Data normalization was performed against the maximum activity index per cage of each disease phase. However, normalized data did not accurately reflect the reduced locomotor capacity experienced by EAE animals (Supplementary Fig. 1). Therefore, raw and unsmoothed data were used for all other data visualization and statistical comparisons. Daily activity index comparison between EAE and sham control mice confirmed the trends observed, showing significant activity reduction in the late-onset (i.e. 13-15dpi) and peak stage of disease (Fig. 1D; two-way RM ANOVA; **24hr**: interaction [activity index x dpi]: F(3.166, 28.49) = 7.291, p = 0.0008). Comparison of daily activity by light phase revealed a larger reduction in activity during the dark/active phase than the light/resting phase (Fig. 1D; two-way RM ANOVA; interaction [activity index x dpi]: **ZT0-12**: F(4.93, 44.37) = 2.949, p = 0.0225; **ZT12-24**: F(3.321, 29.89) = 6.893, p = 0.0008).

**Figure 1.**
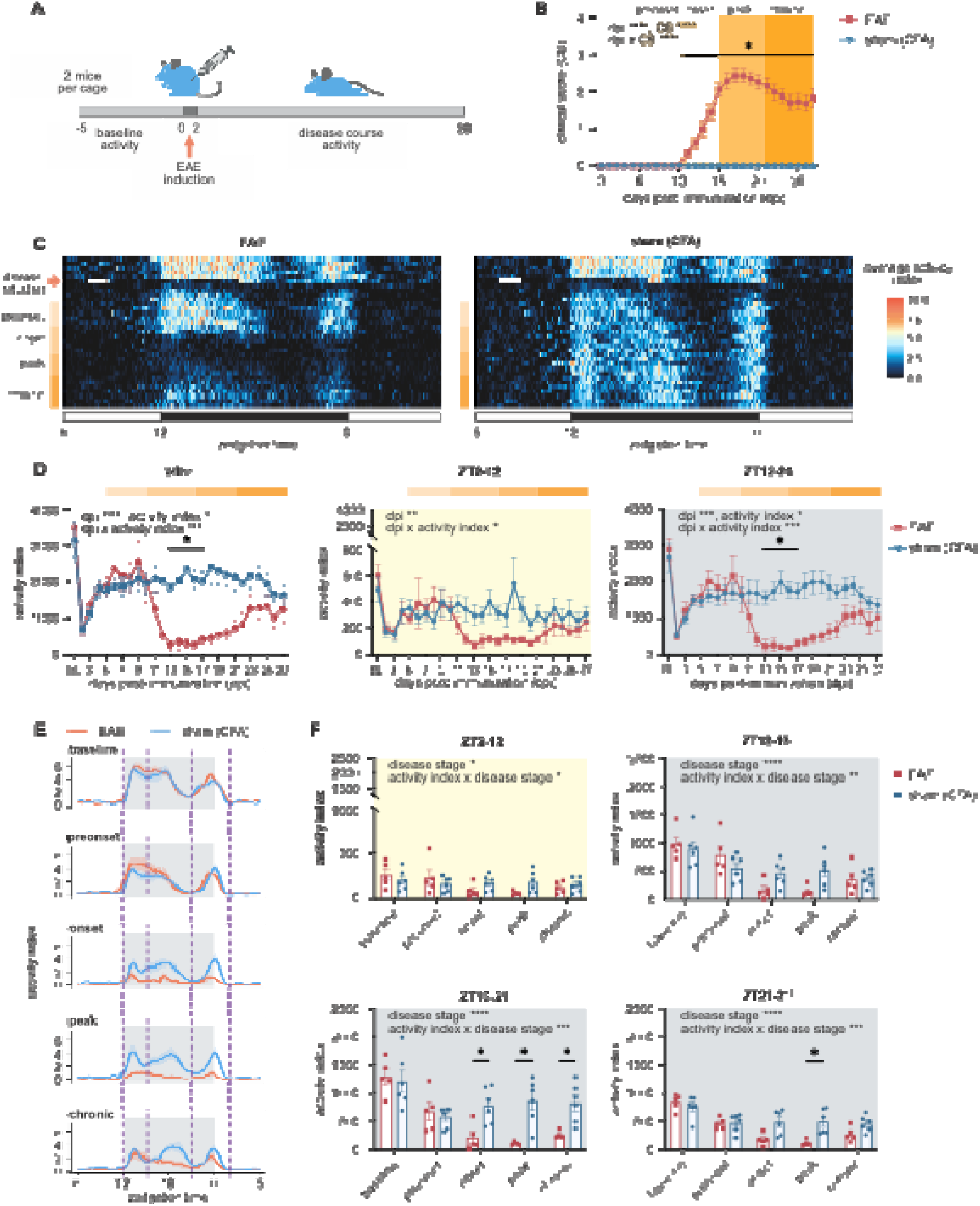
Time-of-day dependent locomotor activity reduction in EAE. **A)** Experiment setup and timeline. **B)** Average clinical score profile of EAE and sham mice and disease stages used for analysis (n = 10-12 mice/group). **C)** Heatmaps of activity index for both EAE and sham mice. **D)** Day-to-day activity index comparison between EAE and sham mice showing 24-hr, resting phase and active phase comparisons. **E)** Activity index distribution over 24 hours in each disease stage with dotted lines showing activity period division based on activity patterns. **F)** Statistical comparison of activity in each period across disease stages. All data analyzed with two-way RM ANOVA with post-hoc Bonferroni’s test; n = 5-6 cages/group, 2 animals/cage. ∗p < 0.05, ∗∗p < 0.01, ∗∗∗p < 0.001, ∗∗∗∗p < 0.0001; all data are represented as mean ± SEM.

Previous studies have demonstrated distinct periods of high activity in the dark phase of rodents (Bains et al., 2018, Usiyevich et al., 2025), corresponding to behaviours such as feeding, grooming and exploring (Hsieh et al., 2024, Istudor et al., 2025). We therefore sought to investigate if the changes observed could be isolated to certain activity periods. The distribution of activity index over 24 hours was plotted for each disease stage; alignment of activity distribution across disease stages led to the identification of four distinct activity periods: ZT2-12, ZT12-15, ZT15-21 and ZT21-2^+1^ (ZT2 of the following day; Fig. 1E). Statistical comparison in each activity period showed significant interactions across all activity periods (Fig. 1F). Activity in the ZT15-21 period showed minimal recovery with improvements in motor deficits, while activity in other periods decreased in the onset and peak phase and recovered in the chronic phase (Fig. 1F; two-way RM ANOVA; interaction [activity index x disease stage]: **ZT2-12**: F(1.907, 17.17) = 4.173, p = 0.0349; **ZT12-15**: F(2.331, 20.98) = 5.679, p = 0.0083; **ZT15-21**: F(2.337, 21.03) = 10.32, p = 0.0005; **ZT21-2^+1^**: F(2.763, 24.87) = 9.115, p = 0.0004). Therefore, our analysis identified a decrease in locomotor activity of EAE mice concentrated during the dark/active phase, with activity reduction lasting from onset to chronic stage, particularly between ZT15-21.

### 3.2 EAE pathology causes activity fragmentation and impairs long-term activity maintenance

An important metric of home-cage activity is the length of activity bouts, which serves as an indicator of activity fragmentation and maintenance (Piilgaard et al., 2023). Activity bouts in each disease stage were categorized based on bout lengths that were divided into bins of 1 minute, 2-5 minutes, 6-10 minutes, 11-20 minutes, 21-40 minutes and >40 minutes duration, as previously described (Piilgaard et al., 2023, Lim et al., 2013). Although previous studies using activity analysis correlated inactive periods of >40 seconds to sleep based on video recording evidence (Pack et al., 2007), our study considered such periods as “rest” to account for the potential immobility caused by motor deficits. Thus, sleep is not included as a measure using these metrics. Stacked area plotting was used to represent normalized activity bout distribution, with the duration of each bin as a percentage of overall activity duration in each disease stage. Activity distribution of sham mice was stable across disease stages, whereas those of diseased mice showed pronounced fluctuations (Fig. 2A). Statistical analysis revealed a significant decrease in the percentage of >40-minute activity bout duration in the onset, peak and chronic stages of disease (Fig. 2B), which suggests a long-term reduction in activity maintenance. Activity fragmentation was most apparent in the peak stage of disease, marked by significant increases in shorter bouts of activity (e.g., 1-minute, 2-5-minute, and 6-10-minute) with a simultaneous decrease in >40-minute activity bouts (Fig. 2B; two-way RM ANOVA; interaction [% total duration x disease stage]: **Baseline**: F(1.273, 11.46) = 0.1728, p = 0.7444; **Preonset**: F(1.435, 12.91) = 0.4180, p = 0.6014; **Onset**: F(2.178, 19.60) = 7.191, p = 0.0038; **Peak**: F(2.126, 19.13) = 28.99; p < 0.0001; **Chronic**: F(2.202, 19.82) = 7.324, p = 0.0034).

**Figure 2.**
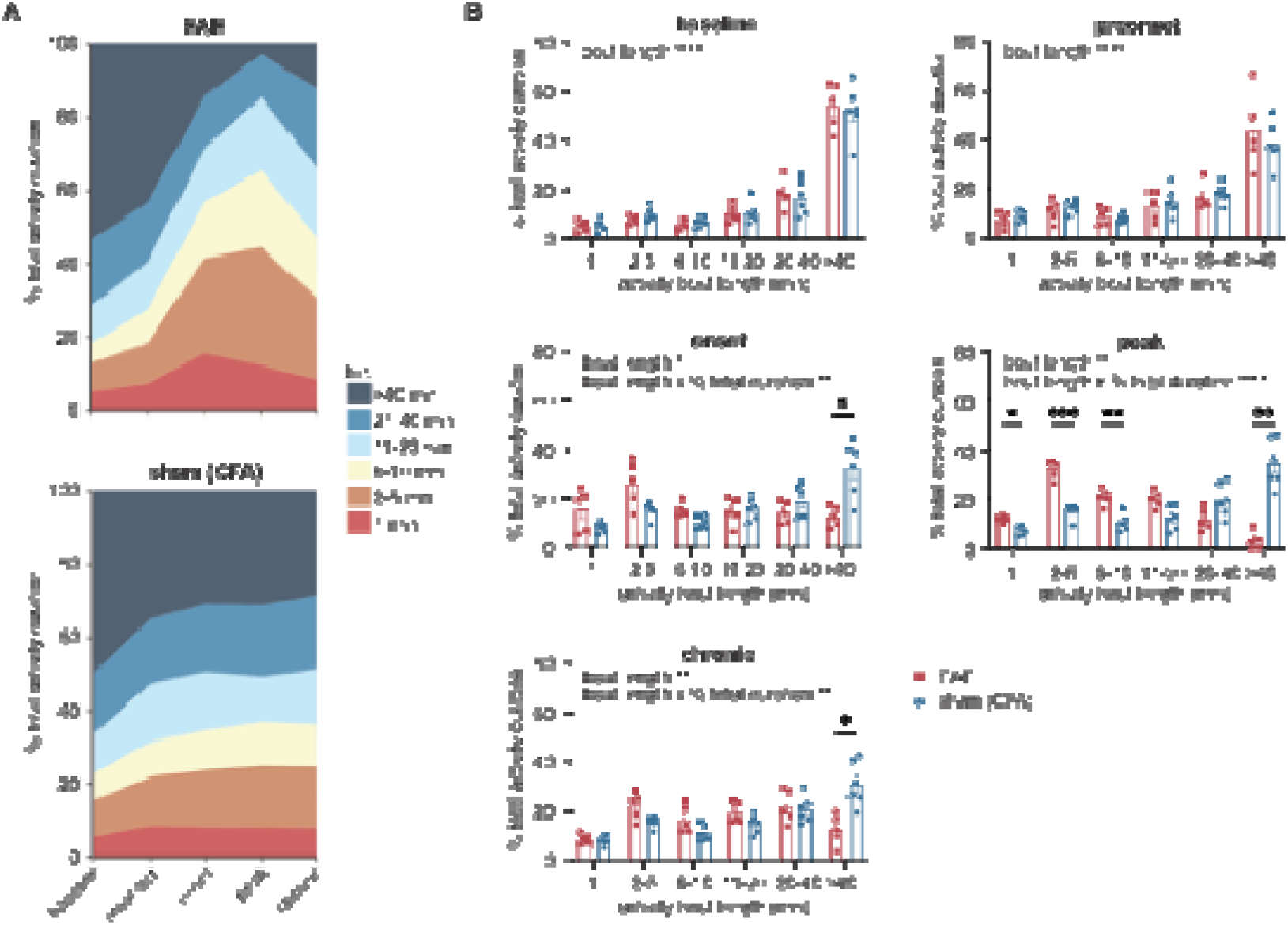
EAE mice exhibit activity fragmentation over the course of disease. **A)** Stacked area plots of changes in activity bout duration in each bin as a percentage of total activity duration in each disease stage. **B)** Statistical comparison of different activity bouts between EAE and sham mice in each disease stage. Two-way RM ANOVA with post-hoc Bonferroni’s test, n = 5-6 cages/group, 2 animals/cage; ∗p < 0.05, ∗∗p < 0.01, ∗∗∗p < 0.001, ∗∗∗∗p < 0.0001; all data are represented as mean ± SEM.

### 3.3 Motor deficit is not the sole contributor to impaired activity maintenance in EAE

We next sought to investigate the correlation between progression of motor deficits and activity maintenance. Dual-axis plots of daily activity bout frequency and clinical score showed differential impact by EAE disease progression on activity bouts of various lengths (Fig. 3A). The same trend was found in other daily activity bout measures, including activity bout index and duration, while daily bout frequency showed the least variability and random fluctuations (Supplementary Fig. 2-6). Diseased mice also showed different frequencies of long activity bouts in the onset and chronic stages that cannot be fully explained by motor deficits. We compared activity bout frequency on consecutive days between the onset and chronic stages of disease (Supplementary Fig. 7). Paired t-tests showed a significantly lower frequency of longer bouts in the onset stage of disease, despite milder motor deficits, relative to the chronic stage when clinical scores were higher (Fig. 3B; Paired t-test; **1 minute**: t(4) = 0.8108, p = 0.4630; **2-5 minute**: t(4) = 0.4047, p = 0.7064; **6-10 minute**: t(4) = 2.683, p = 0.0551; **11-20 minute**: t(4) = 4.913, p = 0.0080; **21-40 minute**: t(4) = 3.325, p = 0.0292; **>40 minute**: t(4) = 3.913, p = 0.0173). Altogether, these results suggest reduced activity maintenance as a result of both motor and non-motor factors over the course of EAE.

**Figure 3.**
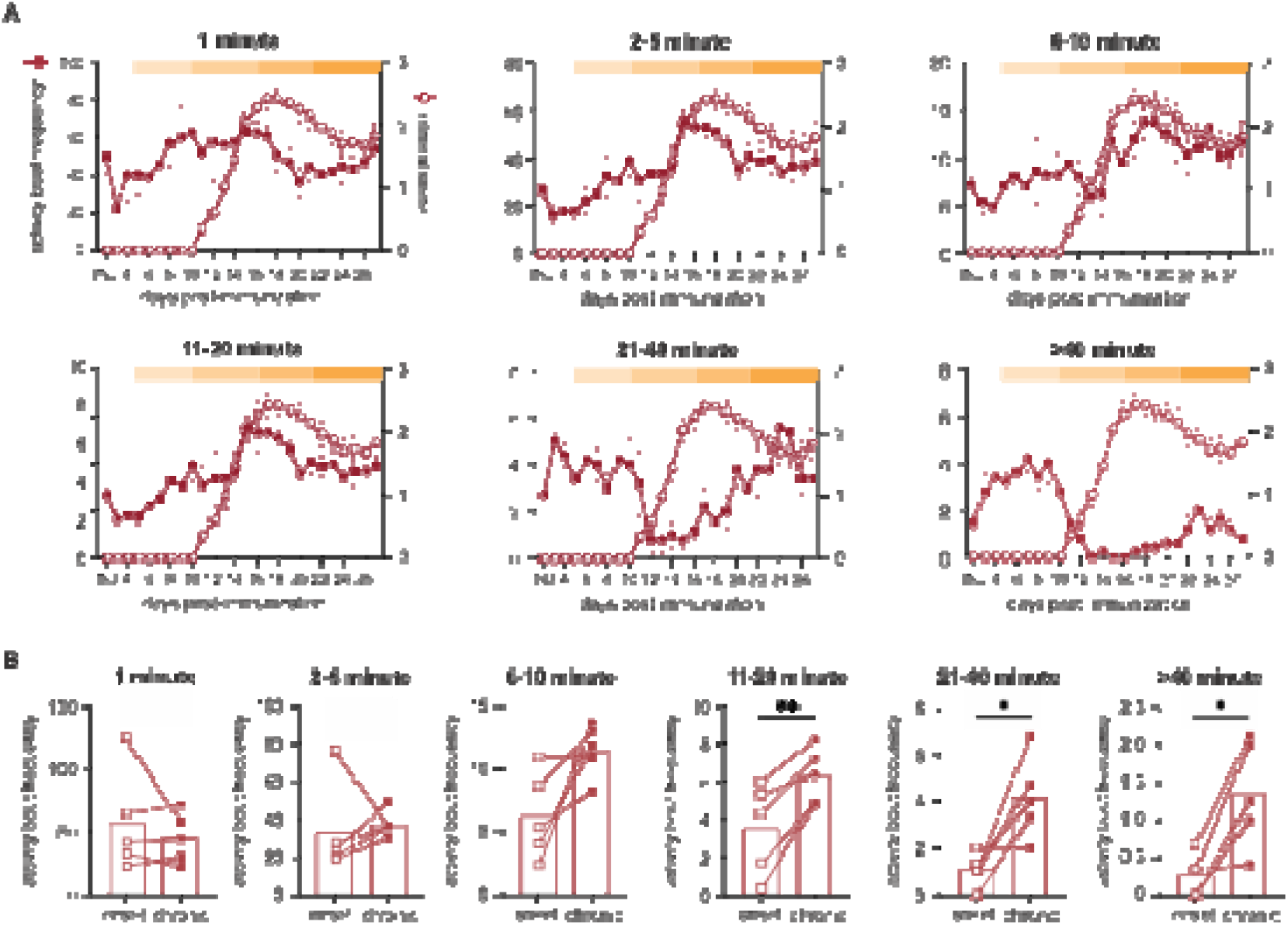
EAE mice show different bout frequency in onset and chronic disease despite similar motor deficits. **A)** Dual-axis plots of correspondence between bout activity and hindlimb paralysis progression; **B)** Daily frequency of activity bouts is compared between days in the onset stage with same or lower clinical scores as days in the chronic stage. Fig. 3B data analyzed with paired Student’s t test, n = 5-6 cages/group, 2 animals/cage; ∗p < 0.05, ∗∗p < 0.01, ∗∗∗p < 0.001, ∗∗∗∗p < 0.0001; all data are represented as mean ± SEM.

### 3.4 Circadian disruption is observed at the peak stage of EAE

Longitudinal data collected using the DVC cages allowed for circadian rhythms analysis throughout the EAE disease course using circadian parameters. Lomb-Scargle analysis was used to examine the periodicity of locomotor activity (Refinetti et al., 2007). Mice exhibited a circadian period of approximately 24 hours at baseline. There was a progressive loss of 24-hour periodicity from onset to peak stage of disease with a partial recovery in the chronic stage, whereas sham controls showed only moderate reduction of periodicity following immunization (Figure 4A). The moderate redution in 24-hour normalized power in sham controls may be due to peripheral inflammation caused by CFA (Lazarevic et al., 2024). Statistical comparison of normalized power in each disease stage showed significantly reduced periodicity in diseased mice compared to sham controls in the peak stage of disease (Figure 4B; two-way RM ANOVA; interaction [normalized power x disease stage]: F(2.662, 23.96) = 7.533, p = 0.0014).

**Figure 4.**
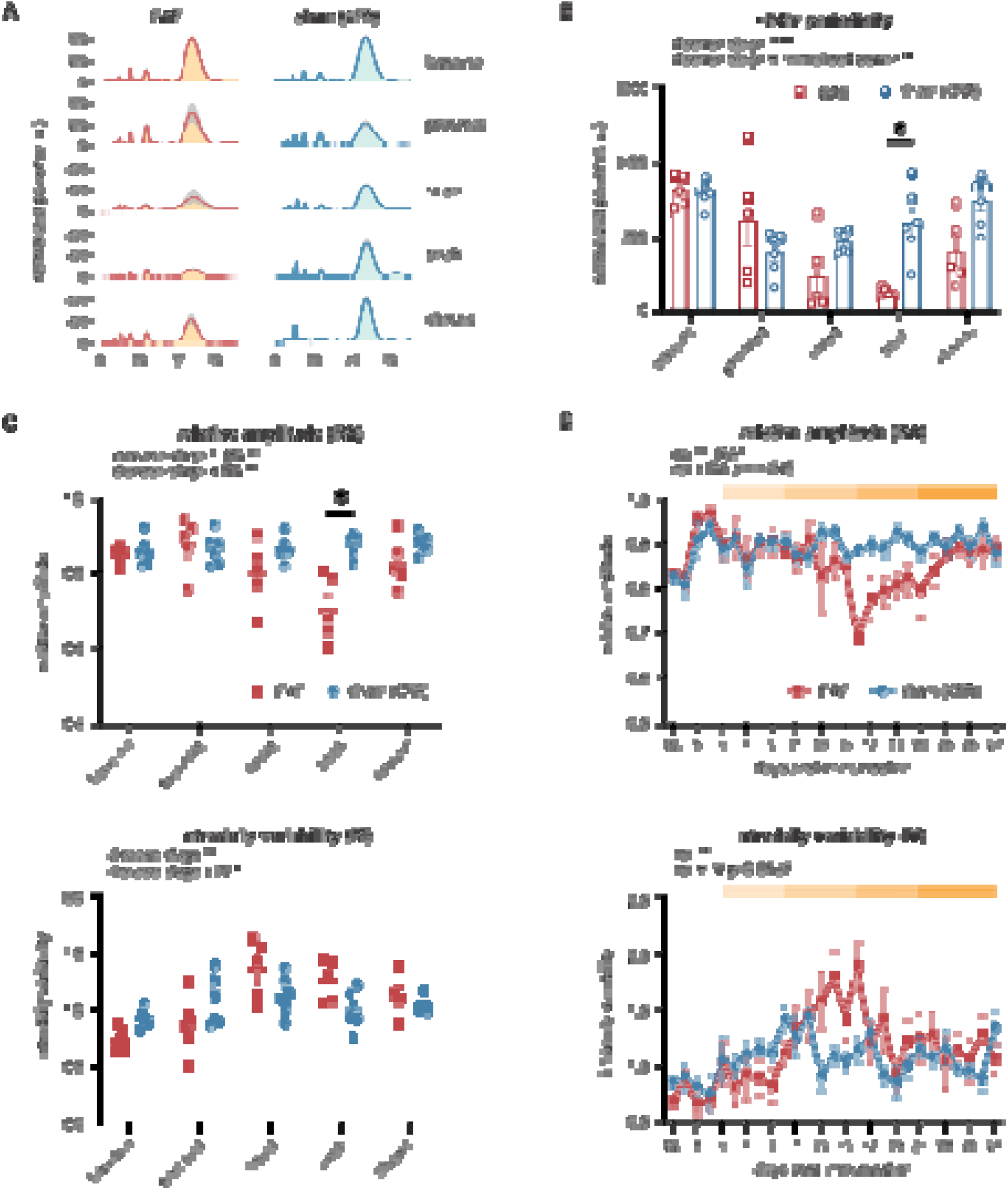
Mice with EAE experience circadian disruption at peak stage of disease. Lombe-Scargle periodogram and non-parametric circadian parameter analysis show changes in circadian rhythmicity after immunization in EAE and sham mice. **A)** Lomb-Scargle periodograms of periodicity across disease stages. **B)** Statistical comparison of ∼24-hour normalized power between EAE and sham mice across disease stages. **C)** Relative amplitude and intradaily variability show circadian disruption at peak disease. **D)** Consecutive 24-hour circadian parameters show near-significant changes in circadian rhythmicity with disease progression. All data analyzed with two-way RM ANOVA with post-hoc Bonferroni’s test; n = 5-6 cages/group, 2 animals/cage; ∗p < 0.05, ∗∗p < 0.01, ∗∗∗p < 0.001, ∗∗∗∗p < 0.0001; all data are represented as mean ± SEM.

We further assessed different aspects of circadian disruption using non-parametric circadian parameters. Comparison by disease stages showed significant changes in relative amplitude and intradaily variability in EAE mice relative to sham controls (Fig. 4C). There was a decrease in relative amplitude at peak stage of disease compared to sham controls, indicating diminished distinction between active and rest phase (Fig. 4C). Intradaily variability, which measures the hour-to-hour rest/active transitions, showed significant interaction but no significant differences between EAE and sham mice in each disease stage by post-hoc analyses (Fig. 4C; two-way RM ANOVA; **RA**: interaction [RA x disease stage], F(2.845, 25.61) = 5.341, p = 0.0060; **IV**: interaction [IV x disease stage], F(2.156, 19.40) = 4.400, p = 0.0244). Daily circadian parameters were used to investigate changes in circadian rhythm through disease progression: both relative amplitude and intradaily variability showed near-significant interaction effects with high between-cage variability (Fig. 4D; two-way RM ANOVA; **RA**: interaction [RA x dpi], F(5.114, 46.03) = 2.344, p = 0.0549; **IV**: interaction [IV x dpi], F(4.531, 40.78) = 2.478, p = 0.0527).

### 3.5 Home-cage activity parameters as indicators of disease progression

Our data generated using the DVC system identified several circadian and locomotor parameters that correlate with disease stages. Changes in these parameters across disease stages were visualized using radar plots (Fig. 5A) and identified that disease onset in EAE mice was marked by increased intradaily variability and 2-5-minute normalized activity bout duration, as well as decreased >40-minute activity bout frequency and normalized power of 24-hour period (Fig. 5A). Peak disease showed similar changes to the onset stage of EAE, with an additional decrease in the relative amplitude not observed at onset. We observed a full recovery in relative amplitude and intradaily variability in the chronic stage, with other parameters partially recovering from peak of disease (Fig. 5A).

**Figure 5.**
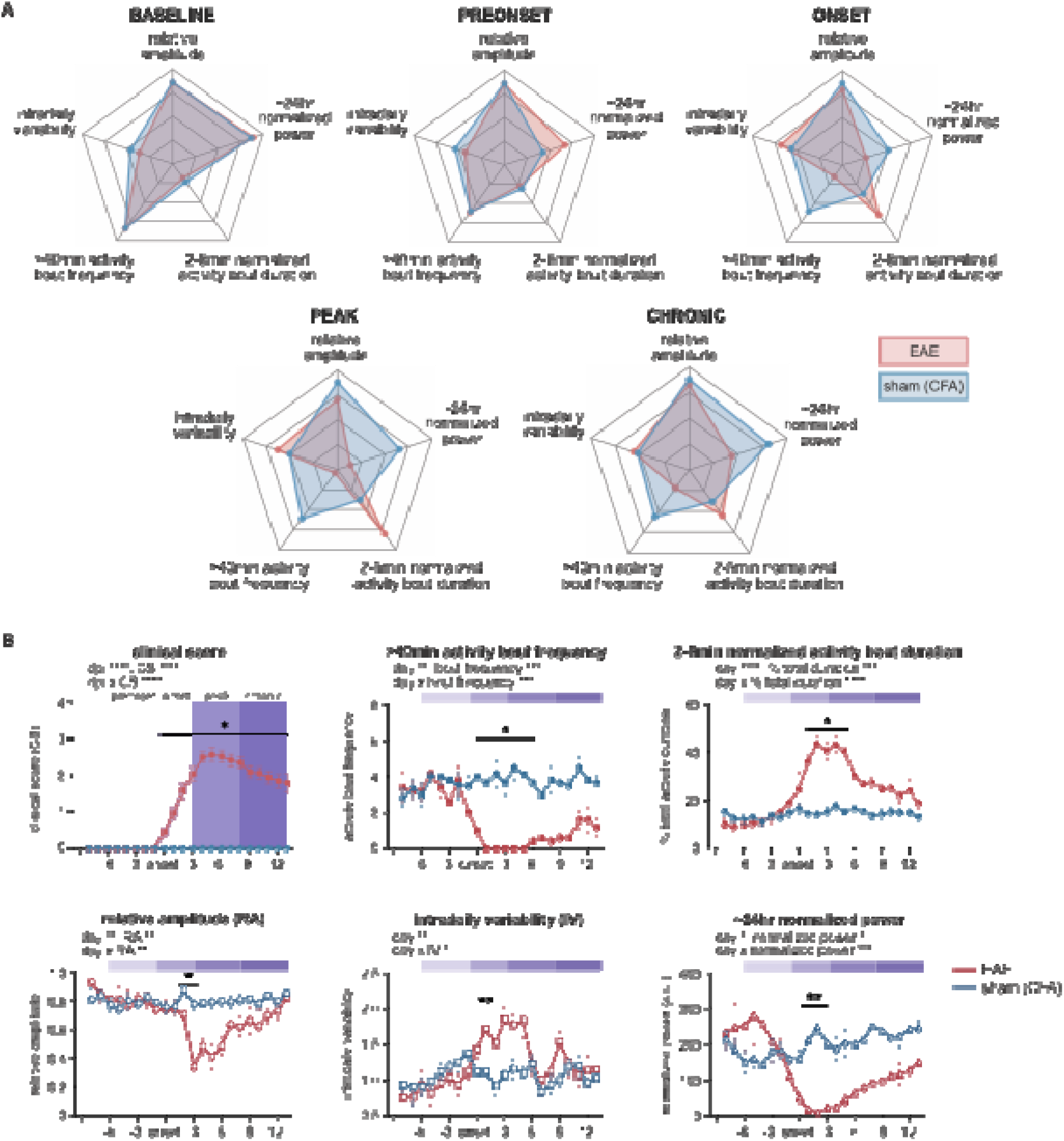
Key circadian and locomotor parameters as indicators of disease progression. **A)** Radar plots demonstrate direction of changes in different parameters across disease stages; **B)** Daily changes in parameters show sensitivity to disease progression in the onset and peak stages of disease. All data analyzed with two-way RM ANOVA with post-hoc Bonferroni’s test; n = 5-6 cages/group, 2 animals/cage; ∗p < 0.05, ∗∗p < 0.01, ∗∗∗p < 0.001, ∗∗∗∗p < 0.0001; all data are represented as mean ± SEM.

While we opted to use average clinical score profiles as a means of determining disease stage in previous analyses, we remained cognizant that EAE typically shows a range of disease onset times between 9 to 14dpi (Constantinescu et al., 2011, Bittner et al., 2014). We therefore sought to explore if activity parameters could be used as indicators of disease *progression*. Daily parameters were aligned between cages, based on the day of disease onset, to assess the sensitivity of parameters to disease progression (Fig. 5B; two-way RM ANOVA; interaction [CS x day]; **Clinical score**: F(2.949, 58.99) = 104.2, p < 0.0001). The most significant change in activity was observed at onset, with all identified activity parameters showing significant changes (Fig. 5B). Frequency of >40-minute activity bouts showed the highest sensitivity to disease onset, with a significant decrease in frequency on the day of disease onset (Fig. 5B; two-way RM ANOVA; interaction [bout frequency x day]; **>40min activity bout frequency**: F(5.617, 50.55) = 5.838, p = 0.0002). Other activity parameters, including 2-5-minute normalized activity bout duration, relative amplitude, intradaily variability and ∼24hr normalized power, showed significant changes 1 day after disease onset (Fig. 5B; two-way RM ANOVA; **2-5min normalized activity bout duration**: interaction [% total duration x day], F(4.891, 44.02) = 11.22, p < 0.0001; **RA**: interaction [RA x day], F(5.449, 49.04) = 4.007, p = 0.0028; **IV**: interaction [IV x day], F(4.502, 40.52) = 3.513, p = 0.0121; **∼24hr normalized power**: interaction [normalized power x day], F(3.510, 31.59) = 6.807, p = 0.0007). Significant changes in >40-minute activity bout frequency and 2-5-minute normalized activity bout duration extended into the peak stage of disease (Fig. 5B). Generally, all identified activity parameters showed significant changes in the onset stage that remained or trended into the peak stage, with chronic stage showing full or partial recovery of activity parameters towards sham levels.

### 3.6 EAE mice with atypical disease show distinct activity phenotype before disease onset

We next explored if home-cage activity could capture atypical disease progression of EAE. A subset of EAE mice in this study developed mild chronic disease with an average clinical score of 1 persisting through the disease course, which differs from the typical relapse-remitting progression pattern (Fig. 6A; two-way RM ANOVA; interaction [CS x dpi]: F(5.258, 81.51) = 15.69, p < 0.0001; One-way ordinary ANOVA; F(2, 81) = 108.8, p < 0.0001). In our experiments, 12 out of 22 or 54.5% of all diseased mice exhibited this atypical disease course, a rate consistent with published literature of EAE model on the C57BL6/J background (Nelson et al., 2024, Mayrhofer et al., 2021); this resulted in 6 out of 11 cages of EAE mice being marked as “atypical EAE” in subsequent analysis. Different disease progression was reflected in body weight changes, where mice with atypical EAE showed mild weight loss compared to their relapse-remitting counterparts (Supplementary Fig. 8). Mice with atypical disease course showed a consistent but moderate decrease in activity index compared to sham controls (Fig. 6B; two-way RM ANOVA; interaction [activity index x dpi]: F(7.067, 49.47) = 5.387, p = 0.0001). We investigated if the decrease in activity index was isolated to specific activity bout lengths.

**Figure 6.**
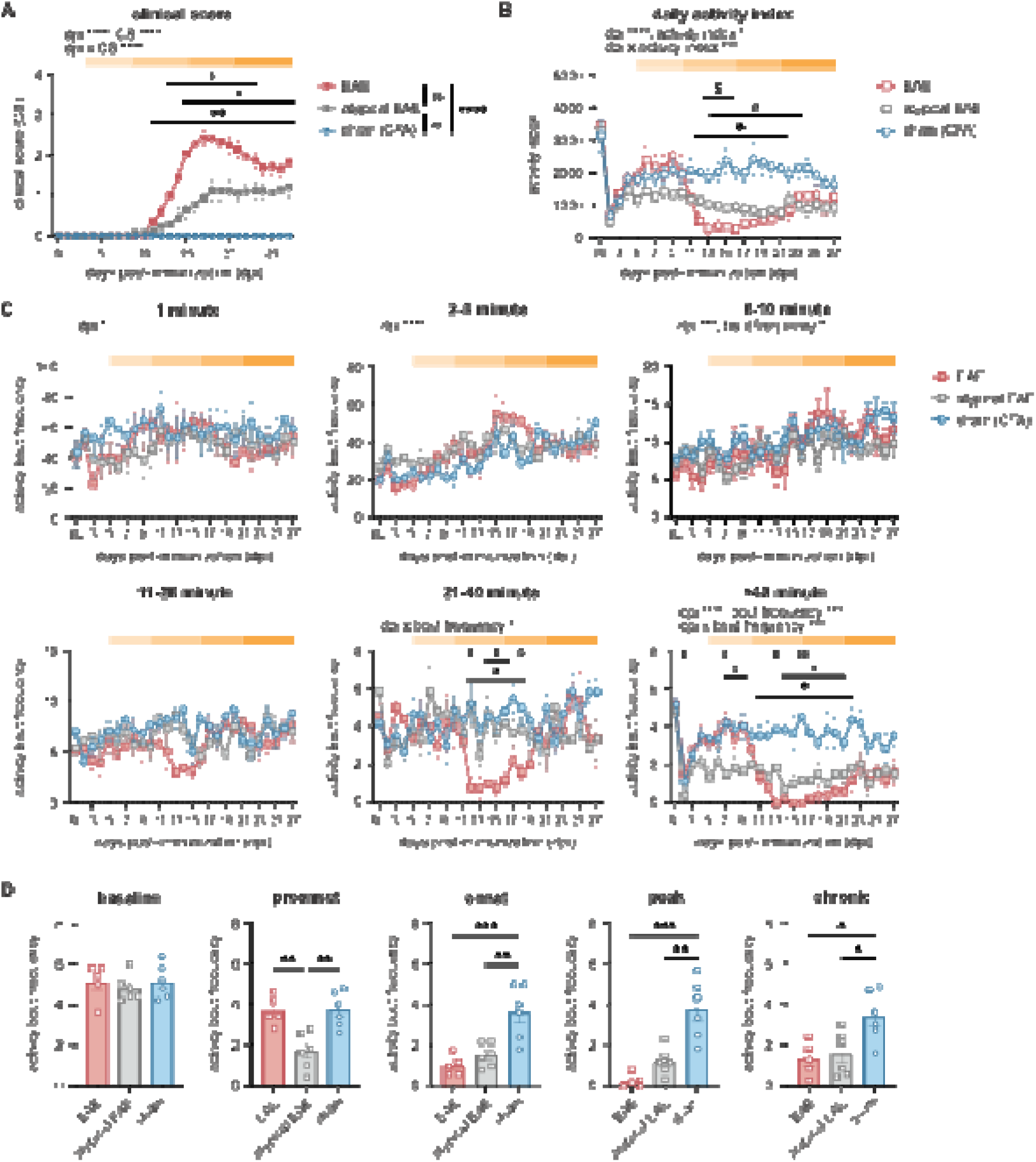
Emergence of a detectable activity phenotype occurs early in atypical EAE disease progression. EAE mice with atypical clinical score exhibit reduced frequency of long bouts before disease onset. **A)** Clinical score profiles of EAE mice, atypical EAE mice and sham mice. **B)** Daily activity index of EAE mice, atypical EAE mice and sham mice. **C)** Daily frequency of various bout lengths of EAE mice, atypical EAE mice and sham mice. **D)** Average daily bout frequency comparison between EAE mice, atypical EAE mice and sham mice per disease stage. Figure 6A between-group comparison and Figure 6D were analyzed with one-way ordinary ANOVA. Figure 6A, B and C were analyzed with two-way RM ANOVA. Tukey’s test was used for all post-hoc comparisons. N = 5-6 cages/group, 2 animals/cage; ∗p < 0.05, ∗∗p < 0.01, ∗∗∗p < 0.001, ∗∗∗∗p < 0.0001; all data are represented as mean ± SEM. ∗ EAE vs sham, ^#^ atypical EAE vs sham, ^$^ EAE vs atypical EAE.

Comparison of daily activity bout frequency showed no difference between sham mice and atypical EAE mice in activity bouts less than 20 minutes (Fig. 6C; two-way RM ANOVA; interaction [bout frequency x dpi]: **1 minute**: F(11.00, 82.49) = 1.083, p = 0.3849; **2-5 minute**: F(12.56, 87.93) = 1.529, p = 0.1253; **6-10 minute**: F(16.25, 113.8) = 0.9091, p = 0.5618; **11-20 minute**: F(15.86, 111.0) = 1.069, p = 0.3932). Frequency of >40-minute activity bouts exhibited similar trends to daily activity index and clinical score, suggesting that the most prominent difference in activity occurred within this bout category (Fig. 6C; two-way RM ANOVA; interaction [bout frequency x dpi]: **21-40 minute**: F(15.45, 108.1) = 2.158, p = 0.0113; **>40 minute**: F(14.08, 98.54) = 3.414, p = 0.0002). Similar effects were also observed in other metrics for >40-minute activity bouts (Supplementary Fig. 2-6). Both EAE and atypical EAE mice experienced a significant reduction in >40-minute activity bout frequency after disease onset compared to sham controls (Fig. 6D; one-way ANOVA; **Onset**: F(2, 14) = 13.19, p = 0.0006; **Peak**: F(2, 14) = 19.19, p < 0.0001; **Chronic**: F(2, 14) = 6.214, p = 0.0117). We also observed a significant reduction in >40-minute activity bout frequency in atypical EAE mice compared to both EAE and sham mice in the pre-onset stage of disease, before activity was affected by motor deficits (Fig. 6D; one-way ordinary ANOVA; **Preonset**: F(2, 14) = 10.80, p = 0.0015). Altogether, the atypical disease progression of EAE mice was evident in their >40-minute activity bout frequency, with distinct activity bout phenotypes both before and after disease onset.

## 4. DISCUSSION

Wearable activity recording sensors are increasingly adopted in human studies and clinical trials to complement the current episodic assessment of MS symptoms (Sparaco et al., 2018, Alexander et al., 2021), yet comparable methods of activity assessment remain to be established in pre-clinical models of disease. In this study, we investigated whether home-cage activity data can be used as an observer-independent tool for monitoring disease progression and spontaneous activity in the EAE model of MS. Our results showed a close correlation between EAE disease progression and home-cage activity, with reductions in locomotor activity counts and long activity bouts as well as disruptions to circadian activity occurring with disease progression. A subset of activity parameters showed correspondence with individual disease stages, indicating their potential as disease stage identifiers. Long bout activity in particular emerged as an attribute that may reflect non-motor symptoms of EAE, showing significant differences in frequency where clinical scores registered similar levels of motor deficits. Overall, our findings indicate that home-cage monitoring reflects high-resolution changes in activity related to EAE disease progression and merits further investigation as a continuous assessment tool for functional outcomes in the EAE model.

Previous studies have shown a different, bimodal activity pattern in the nocturnal phase, with peaks of anticipatory activity during lights-on and lights-off times (Paladino et al., 2013, Pernold et al., 2021). Strain differences have been shown to be a potential cause for slight variations in activity distribution (van Dorp et al., 2024, Loos et al., 2014, Fuochi et al., 2021). Here, EAE mice experienced activity reduction in all activity periods across both diurnal and nocturnal phases as a result of diminishing mobility in their hindlimbs. Comparison between EAE and sham animals during individual activity periods revealed the most severe activity reduction in the mid-dark phase (i.e. ZT15-21). Unlike other periods that showed activity recovery with mobility improvements in the chronic stage of disease, activity reduction in the mid-dark phase showed no meaningful recovery in chronic disease. Mice have been shown to exhibit the highest frequency of food-seeking behavior during the mid-dark phase, suggesting that the activity reduction may be behaviour-specific (Usiyevich et al., 2025). Sham mice in this study showed a decreasing trend in locomotor activity after immunization with saline/CFA emulsion, which is known to cause peripheral inflammation (Lazarevic et al., 2024) and a moderate but long-lasting reduction in locomotor activity compared to the pre-immunization baseline period (Lu et al., 2012). A recent study using wearable sensor-recorded activity showed similar time-specific activity reduction in pwMS, where lower activity between 12:00-22:00 (or the mid-to late-active phase) was most correlated with variations in EDSS scores (Keller et al., 2022). It is worth noting that the time window of activity reduction is likely specific to MS pathophysiology, as different patterns of activity reduction were found in other conditions with autoimmune components such as rheumatoid arthritis (Prioreschi et al., 2013) and Parkinson’s disease (Nagaki et al., 2022). Altogether, activity reduction in specific time-of-day is an emerging observation in both preclinical and clinical activity studies, and could be related to levels of disability and other behaviours or symptoms.

An important outcome of MS is circadian disruption, which often manifests as reduced integrity of active/rest phases, daytime somnolence, and nighttime awakening (Newland et al., 2016, Sakkas et al., 2019). A broad range of potential mechanisms could contribute to this disruption (Brecier et al., 2023, De Somma et al., 2018). For example, pwMS have significantly reduced melatonin levels compared to healthy controls, with the secretion of melatonin triggering sleep (Melamud et al., 2012) and thus further impacting sleep cycles. Inflammation in the CNS also disrupts the hypothalamus-pituitary-adrenal axis, resulting in impaired sleep/wake cycles and, in turn, circadian rhythms (Fitzgerald et al., 2025). Optic neuritis is a common symptom among pwMS that can lessen photosensitivity of the retino-hypothalamic tract that directs environmental lighting as circadian signals to the hypothalamus (Pivovarova-Ramich et al., 2023). Demyelinating lesions to the hypothalamic region are prevalent in MS, which causes altered cortisol levels and interfere with the sleep/wake response (Burfeind et al., 2016, Kantorova et al., 2017). Thus, circadian and sleep/wake cycles can be profoundly disrupted in pwMS. In this study, EAE mice exhibited disrupted circadian activity in the form of reduced 24-hour periodicity in the peak stage, as well as significant changes in relative amplitude and intradaily variability, which agrees with accelerometry studies in pwMS (Salomon et al., 2022, Bou Rjeily et al., 2025). The exact mechanisms contributing to circadian disruption in EAE, however, remain unexplored but are likely to be similar to those observed in humans.

This study also identified long activity bouts, or activity bouts longer than 40 minutes, as an important attribute of home-cage activity in mice with EAE. Daily frequency of long activity bouts overall correlated with progression of clinical scores. However, our results also showed that long activity bouts could be influenced by non-motor factors. The general trend demonstrated by EAE mice in this study is a decrease in long bout frequency with increasing motor deficits as a result of disease progression. However, average daily long bout frequency was lower in the onset of disease, which was selected for lower motor deficits, than the chronic days. This contradictive phenomenon indicates the potential influence of a non-motor factor in long activity bouts or activity maintenance. The onset stage of EAE is characterized by infiltration of peripheral immune cells into the CNS and activation of CNS-resident glia (Borjini et al., 2016), whereas the chronic stage is marked by low grade or resolving inflammation with chronic degeneration (Hampton et al., 2008, Voskuhl and MacKenzie-Graham, 2022). Inflammation, which is a hallmark of EAE disease onset (Brown and Sawchenko, 2007), is a major contributor towards fatigue behaviour – one of the most common and debilitating symptoms of MS. Recent studies on wearable recordings of pwMS showed a correlation between self-reported fatigue levels and reduced activity maintenance (Blikman et al., 2015, Salomon et al., 2022), a pattern consistent with the reduced frequency of long activity bouts in EAE mice observed in this study. Reduced activity maintenance in pwMS has also been correlated with disease progression, which is typically driven by inflammation. The reduced long activity bout frequency observed in the onset stage despite the less severe motor deficits might therefore be a reflection of fatigue-like behaviour as a result of inflammation.

We also sought to determine whether home-cage activity parameters could be used as indicators of EAE disease progression, in an effort to identify an observer-independent method to assess disease outcomes. The current “gold standard” for assessing EAE disease progression is the clinical score scale (Palle et al., 2016), a subjective assessment that is prone to observer bias (Palle et al., 2016). Moreover, since the score is primarily based on motor deficits, pathological progression that manifests as non-motor symptoms (e.g., inflammation-induced fatigue, altered sleep/wake cycles, etc.) are not captured. Previous literature has proposed other measures of disease progression, including advanced dynamic weight bearing (Segal et al., 2021) and horizontal and vertical activity counts (Shahi et al., 2019). Key activity parameters, including >40-minute activity bout frequency, 2-5-minute normalized activity bout duration, relative amplitude, intradaily variability, and normalized power of 24-hour periodicity, showed differential and pronounced changes at onset and peak stages that resolved in the chronic stage, making them potential identifiers for disease progression. Mice with both typical and atypical EAE disease profiles exhibited close alignment of long activity bout frequency with clinical score profiles, suggesting its utility as a sensitive indicator of disease progression. Mice with atypical EAE also showed fewer long activity bouts than sham and EAE mice before the onset of motor deficits, indicating that long activity bouts may be an early indicator of abnormal disease progression. While various factors can contribute to high variability in the EAE model (Constantinescu, 2005), an early behaviour indicator could aid researchers in efficient experiment design and potentially study underlying biological factors that may contribute to this alternative disease profile.

Several limitations exist in this study. Firstly, we only used female mice in the EAE model, though it is representative of the prevalence of MS in females (Bove and Chitnis, 2013). Importantly, recent studies have shown sex differences in the fatigue-related symptoms of MS; thus, future studies will include male mice for this type of analysis to explore sex differences in fatigue-like symptoms of EAE. Secondly, we focused on the assessment of activity but did not investigate sleep patterns in EAE mice. Since the DVC system utilizes electromagnetic zone crossing as proxy for activity, it cannot differentiate between stationary rest and sleep in an animal model with motor impairment, contributing to the large variability in the rest patterns captured (Supplementary Fig. 9-12). Sleep structure is most effectively investigated using electroencephalogram or electromyogram, which require extracranial or muscle implants on single-housed mice (Mang et al., 2014). Other non-invasive home-cage methods may be used, such as piezoelectric sensor-based monitoring, which captures autonomic movements such as breathing and heartbeats that can be used to infer sleep states (Carreno-Munoz et al., 2022, Mang et al., 2014). Moreover, we housed two animals per cage in this study, to reduce the impact of stress and the documented depressive behaviour in the EAE model (Pollak et al., 2002, Peres et al., 2023). This could confound activity patterns where EAE mice in the same cage have different disease severity or onset times. In an effort to reduce the impact this could have on our results, we only included cages where clinical score profiles are similar between animals, with less than 0.5 clinical score difference between cage mates. Although we observed lower frequency of long activity bouts during the pre-onset stage in atypical EAE mice, this behavioural phenotype has not been corroborated by pathological evidence. Studies have demonstrated a cascade of immune responses in the periphery (Barthelmes et al., 2016) and blood-brain barrier (Alvarez et al., 2015, Davalos et al., 2012, Davalos et al., 2014) before EAE disease onset. During immunization (0-2dpi), mice with atypical EAE showed greater loss of body weight loss and reduced long activity bouts compared to EAE mice. Mice with atypical EAE only had lower body weights when compared to typical EAE mice prior to disease onset (5-9dpi).. The differences observed during immunization and pre-onset stage mayindicate different immune response cascades causing variability in disease progression. Overall, there is sparse evidence connecting these pre-onset events to variability in disease progression. As research interest in the prodromal stage of MS continues to grow (Makhani and Tremlett, 2021), future studies may shed light on how variations in molecular mechanisms during the pre-onset stage could influence disease severity and progression.

This work constitutes the first demonstration of home-cage monitoring to track disease progression and spontaneous activity in the EAE model of MS, laying the groundwork for capturing non-motor symptoms in preclinical models of disease. Screening of generated activity parameters revealed a preliminary batch of disease stage identifiers to expand the repertoire of observer-independent measures for EAE disease progression. This study also presented the first activity-level observation indicative of abnormal disease progression before the onset of disease. As animal housing systems become more widely adopted for colony management, observer-independent measures of home-cage activity could enable high-throughput activity data collection and improve the reproducibility and translational potential of EAE research.

## Availability of data and materials

Data supporting the conclusion of this article are available upon request to the corresponding author.

## Author contribution

**Vina W. Li:** Writing – review & editing, Writing – original draft, Methodology, Investigation, Project administration, Data curation, Software, Formal analysis, Visualization, Conceptualization. **Julia P. Segal:** Methodology, Investigation. **Nader Ghasemlou:** Writing – review & editing, Resources, Methodology, Project administration, Supervision, Funding acquisition, Conceptualization.

## SUPPLEMENTAL DATA

**Supplementary Figure 1.**
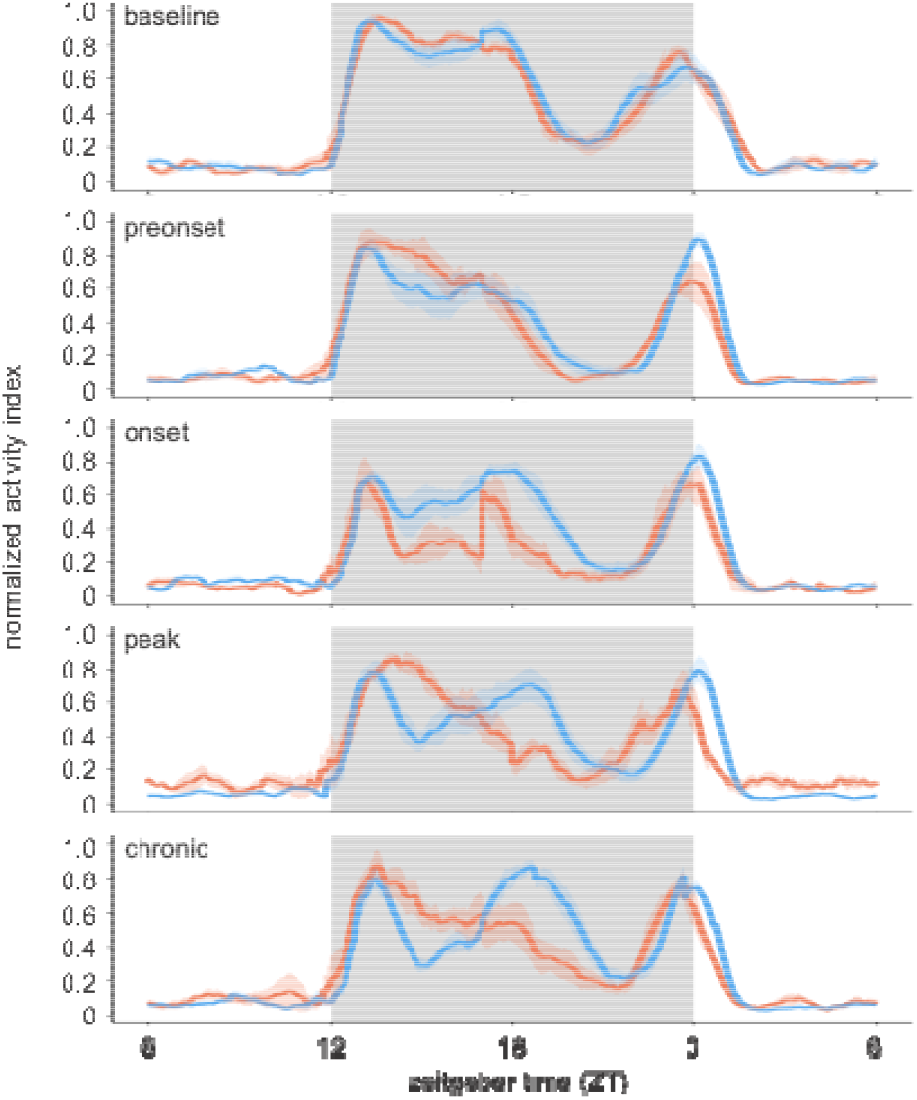
Normalized activity distribution over 24 hours in each disease stage.

**Supplementary Figure 2.**
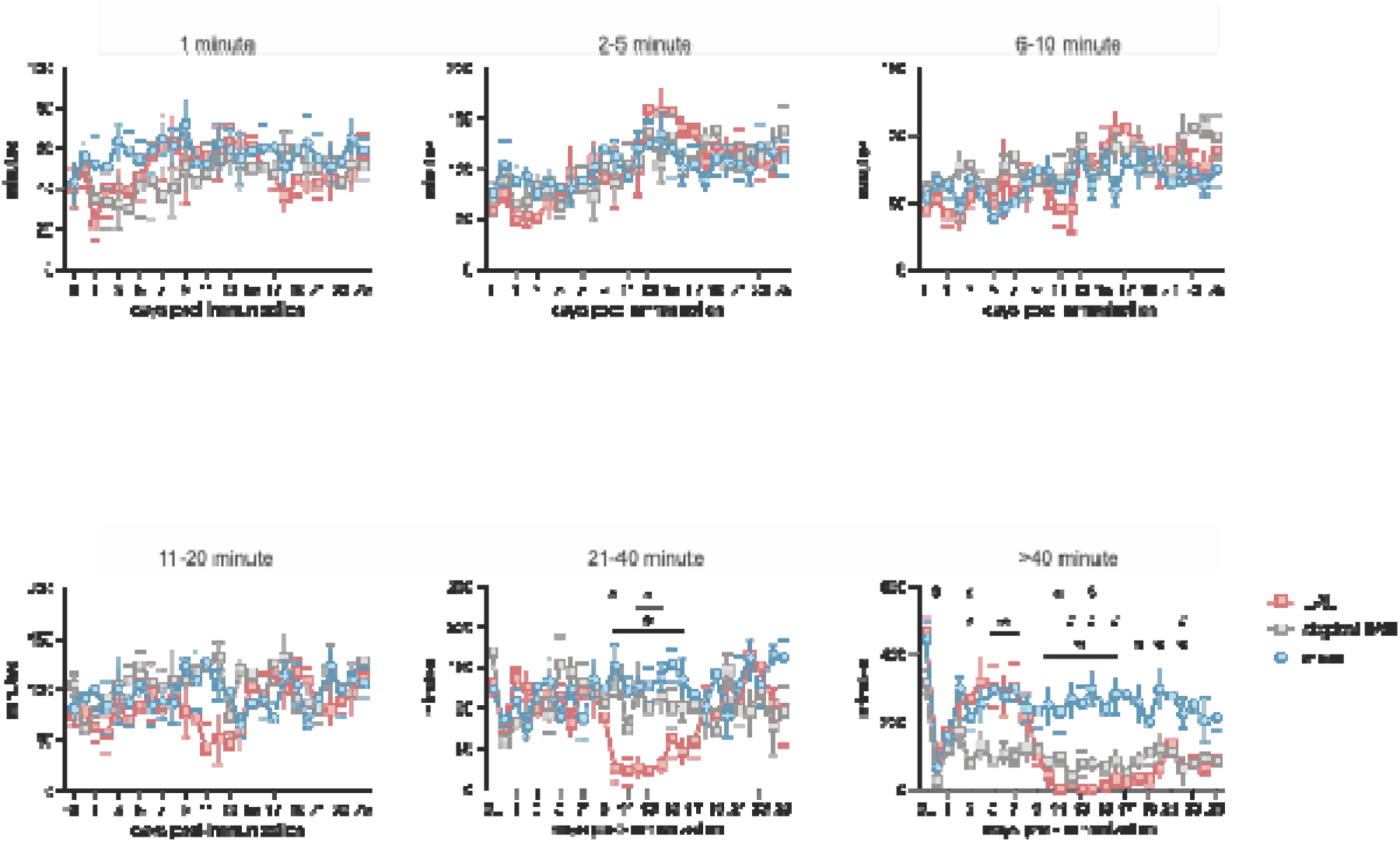
Daily total duration of activity bouts at various lengths in EAE, atypical EAE and sham mice. All data analyzed with two-way RM ANOVA with post-hoc Tukey’s test; n = 5-6 cages/group, 2 animals/cage; ∗p < 0.05, ∗∗p < 0.01, ∗∗∗p < 0.001, ∗∗∗∗p < 0.0001; all data are represented as mean ± SEM. ∗ EAE vs sham, ^#^ atypical EAE vs sham, ^$^ EAE vs atypical EAE.

**Supplementary Figure 3.**
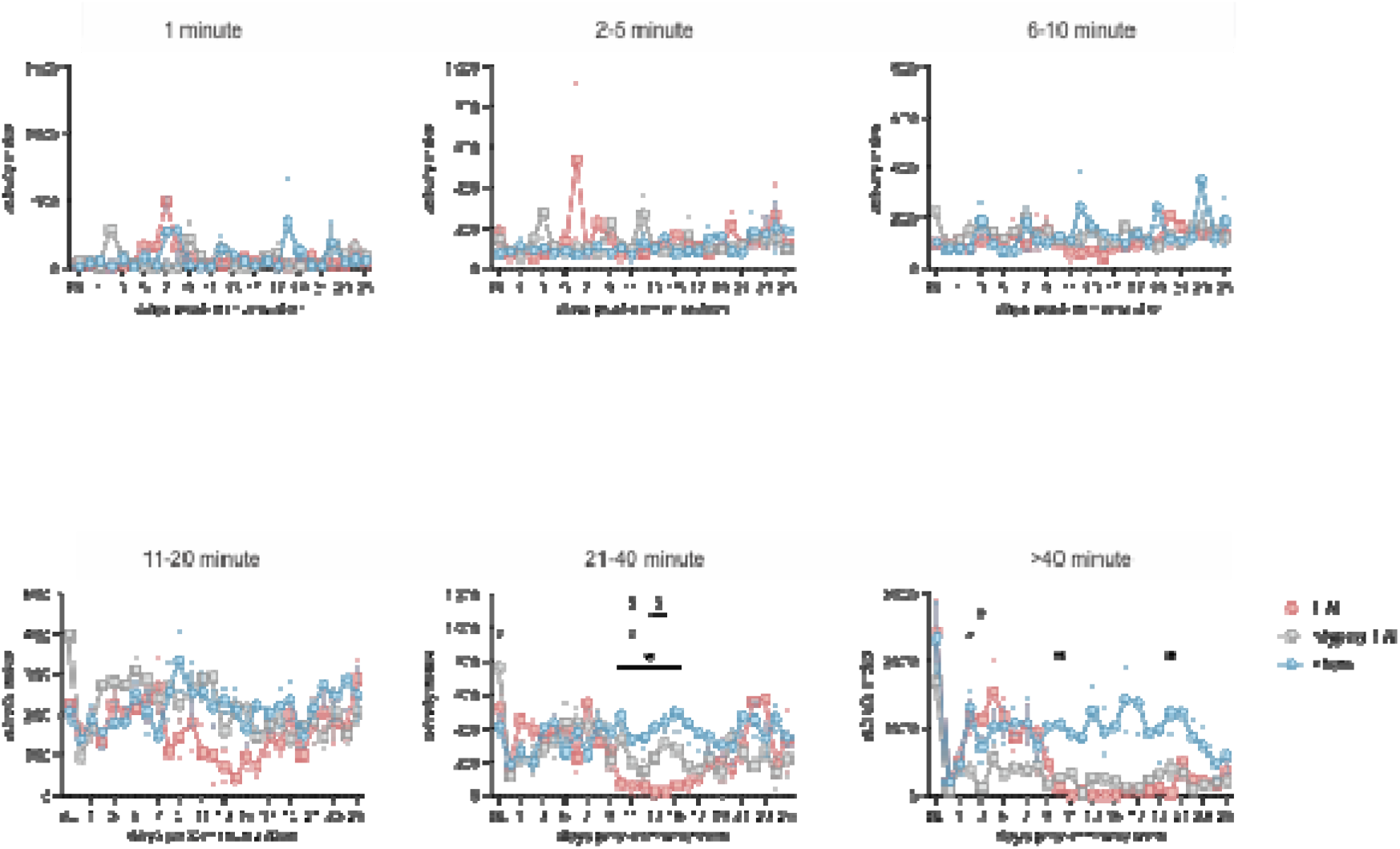
Daily total index of activity bouts at various lengths in EAE, atypical EAE and sham mice. All data analyzed with two-way RM ANOVA with post-hoc Tukey’s test; n = 5-6 cages/group, 2 animals/cage; ∗p < 0.05, ∗∗p < 0.01, ∗∗∗p < 0.001, ∗∗∗∗p < 0.0001; all data are represented as mean ± SEM. ∗ EAE vs sham, ^#^ atypical EAE vs sham, ^$^ EAE vs atypical EAE.

**Supplementary Figure 4.**
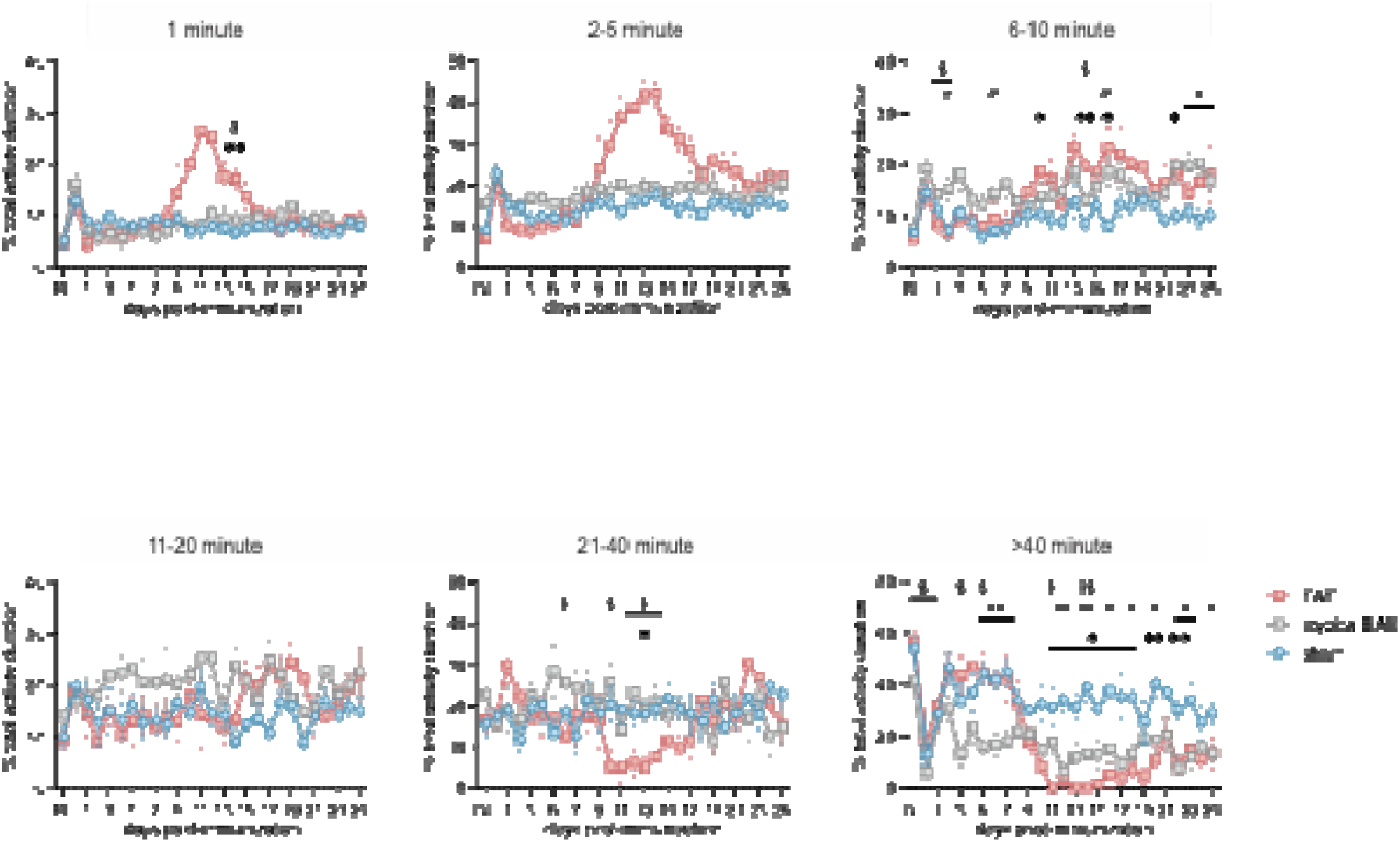
Daily normalized duration of activity bouts at various lengths in EAE, atypical EAE and sham mice. All data analyzed with two-way RM ANOVA with post-hoc Tukey’s test; n = 5-6 cages/group, 2 animals/cage; ∗p < 0.05, ∗∗p < 0.01, ∗∗∗p < 0.001, ∗∗∗∗p < 0.0001; all data are represented as mean ± SEM. ∗ EAE vs sham, ^#^ atypical EAE vs sham, ^$^ EAE vs atypical EAE.

**Supplementary Figure 5.**
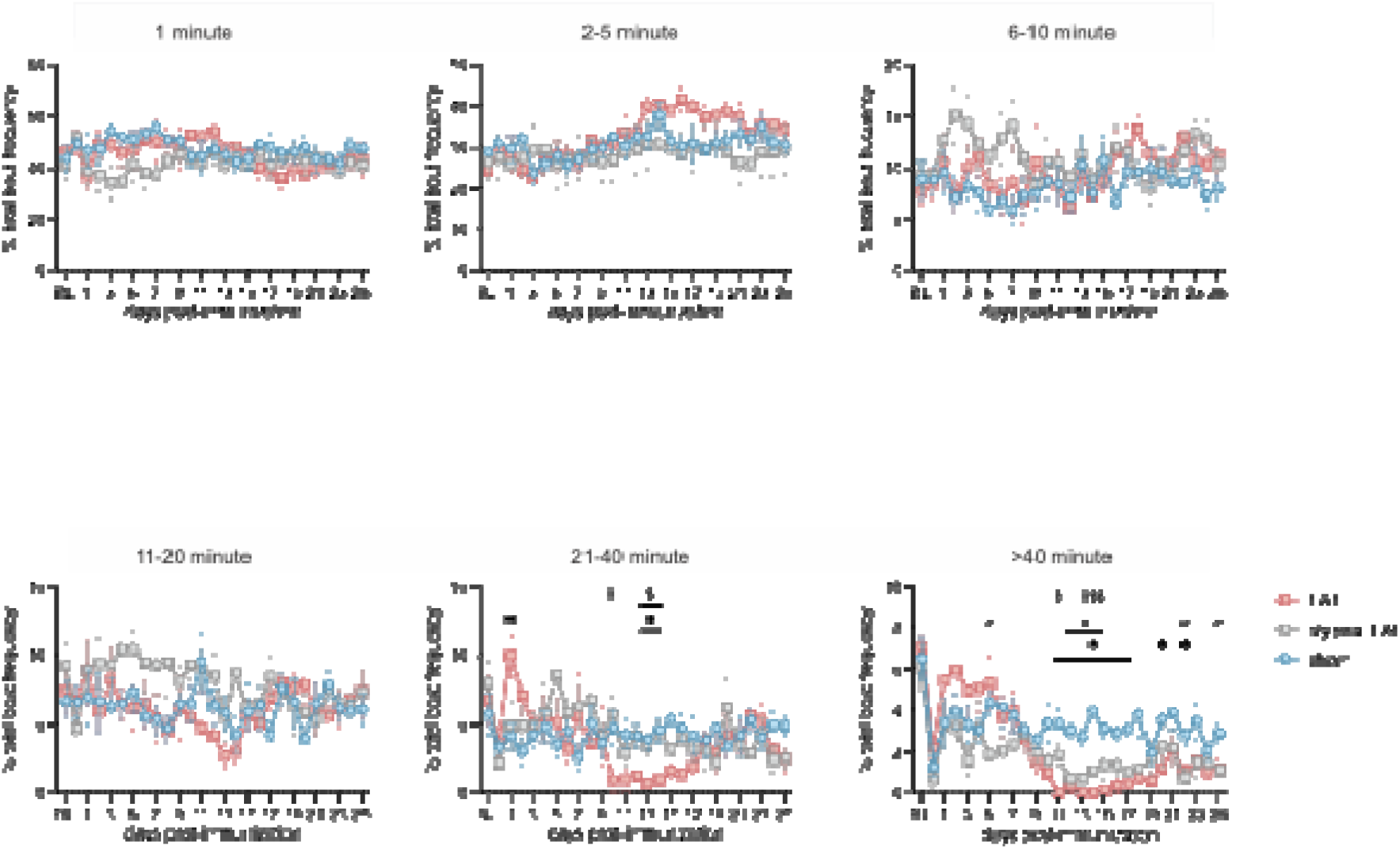
Daily normalized frequency of activity bouts at various lengths in EAE, atypical EAE and sham mice. All data analyzed with two-way RM ANOVA with post-hoc Tukey’s test; n = 5-6 cages/group, 2 animals/cage; ∗p < 0.05, ∗∗p < 0.01, ∗∗∗p < 0.001, ∗∗∗∗p < 0.0001; all data are represented as mean ± SEM. ∗ EAE vs sham, ^#^ atypical EAE vs sham, ^$^ EAE vs atypical EAE.

**Supplementary Figure 6.**
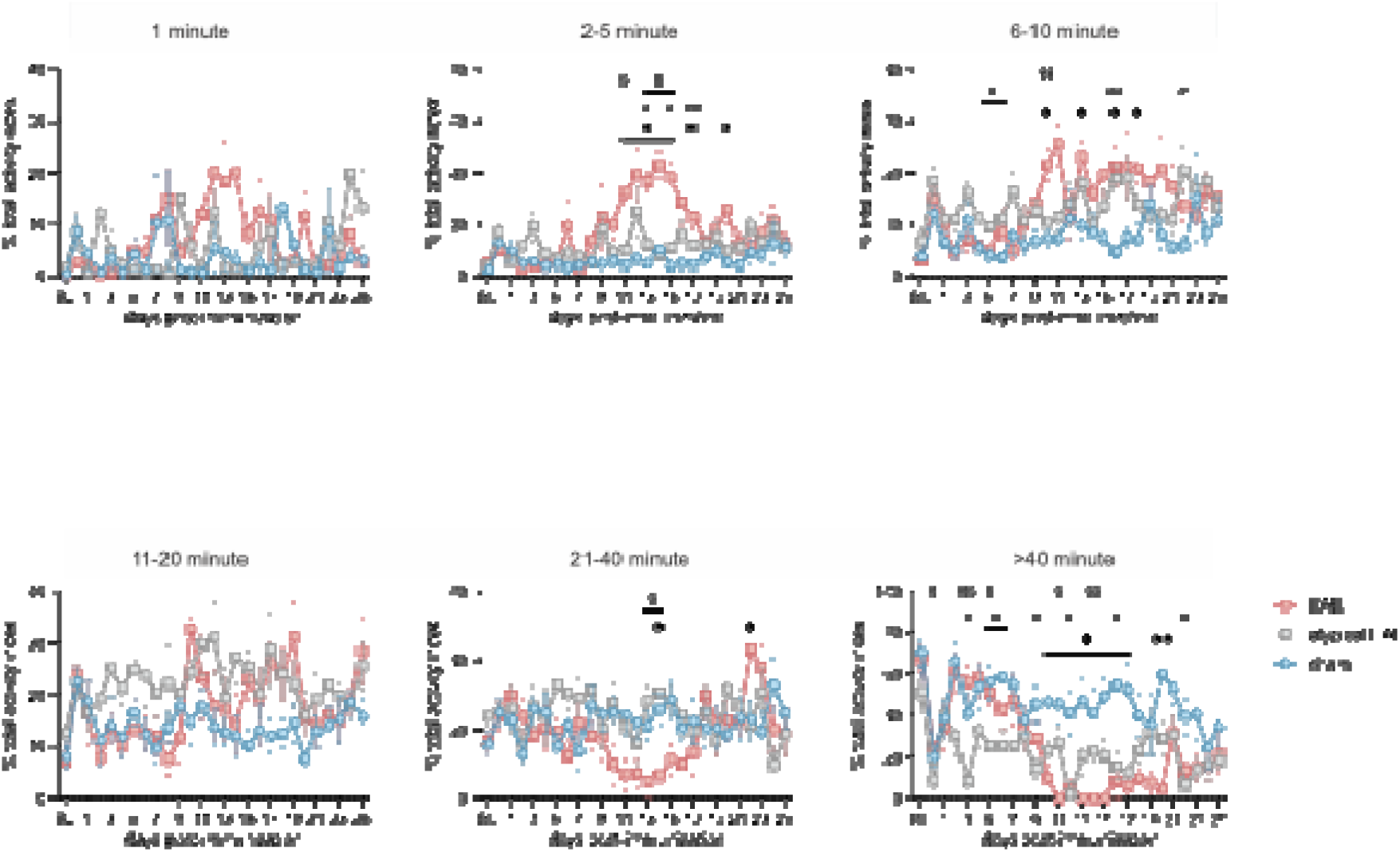
Daily normalized index of activity bouts at various lengths in EAE, atypical EAE and sham mice. All data analyzed with two-way RM ANOVA with post-hoc Tukey’s test; n = 5-6 cages/group, 2 animals/cage; ∗p < 0.05, ∗∗p < 0.01, ∗∗∗p < 0.001, ∗∗∗∗p < 0.0001; all data are represented as mean ± SEM. ∗ EAE vs sham, ^#^ atypical EAE vs sham, ^$^ EAE vs atypical EAE.

**Supplementary Figure 7.**
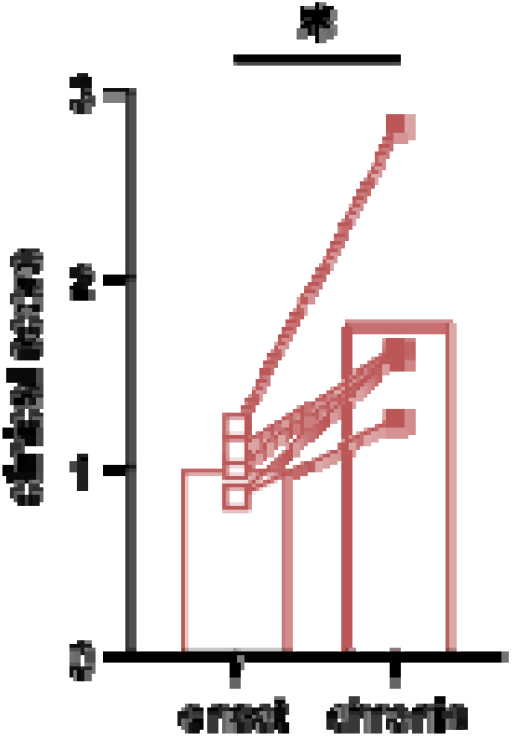
Average clinical score of EAE mice on the days in onset and chronic disease stage selected for paired comparison of daily frequency of activity bouts. Data analyzed with paired Student’s t test; n = 5-6 cages/group, 2 animals/cage; ∗p < 0.05, ∗∗p < 0.01, ∗∗∗p < 0.001, ∗∗∗∗p < 0.0001.

**Supplementary Figure 8.**
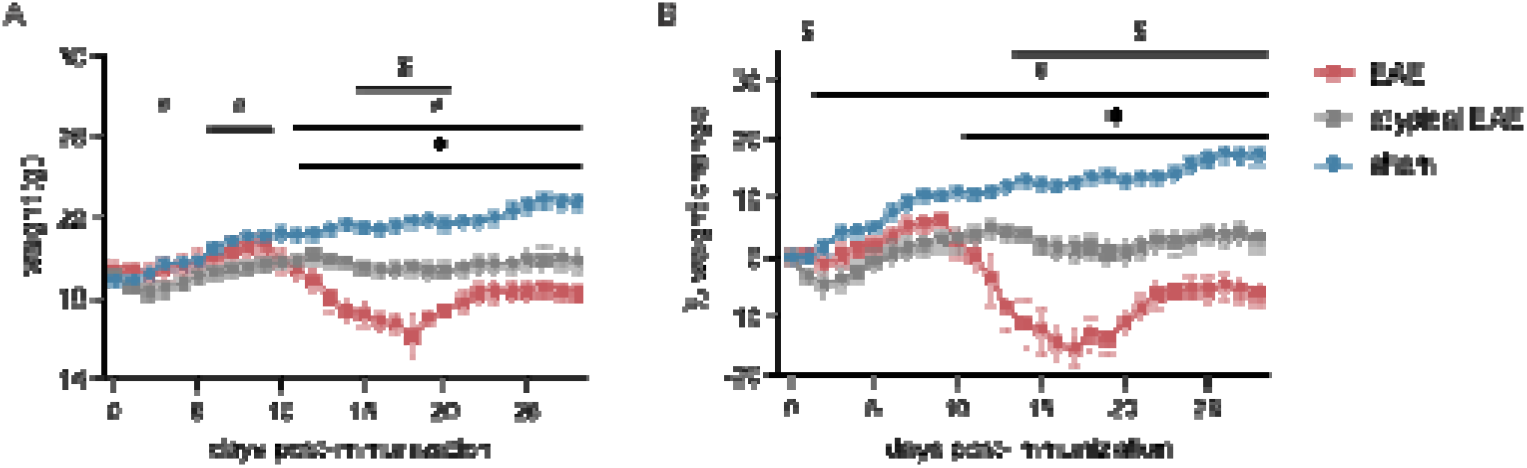
Daily body weights of mice with EAE, atypical EAE and sham disease course. All data analyzed with two-way RM ANOVA with post-hoc Tukey’s test; n = 5-6 cages/group, 2 animals/cage; ∗p < 0.05, ∗∗p < 0.01, ∗∗∗p < 0.001, ∗∗∗∗p < 0.0001; all data are represented as mean ± SEM. ∗ EAE vs sham, ^#^ atypical EAE vs sham, ^$^ EAE vs atypical EAE.

**Supplementary Figure 9.**
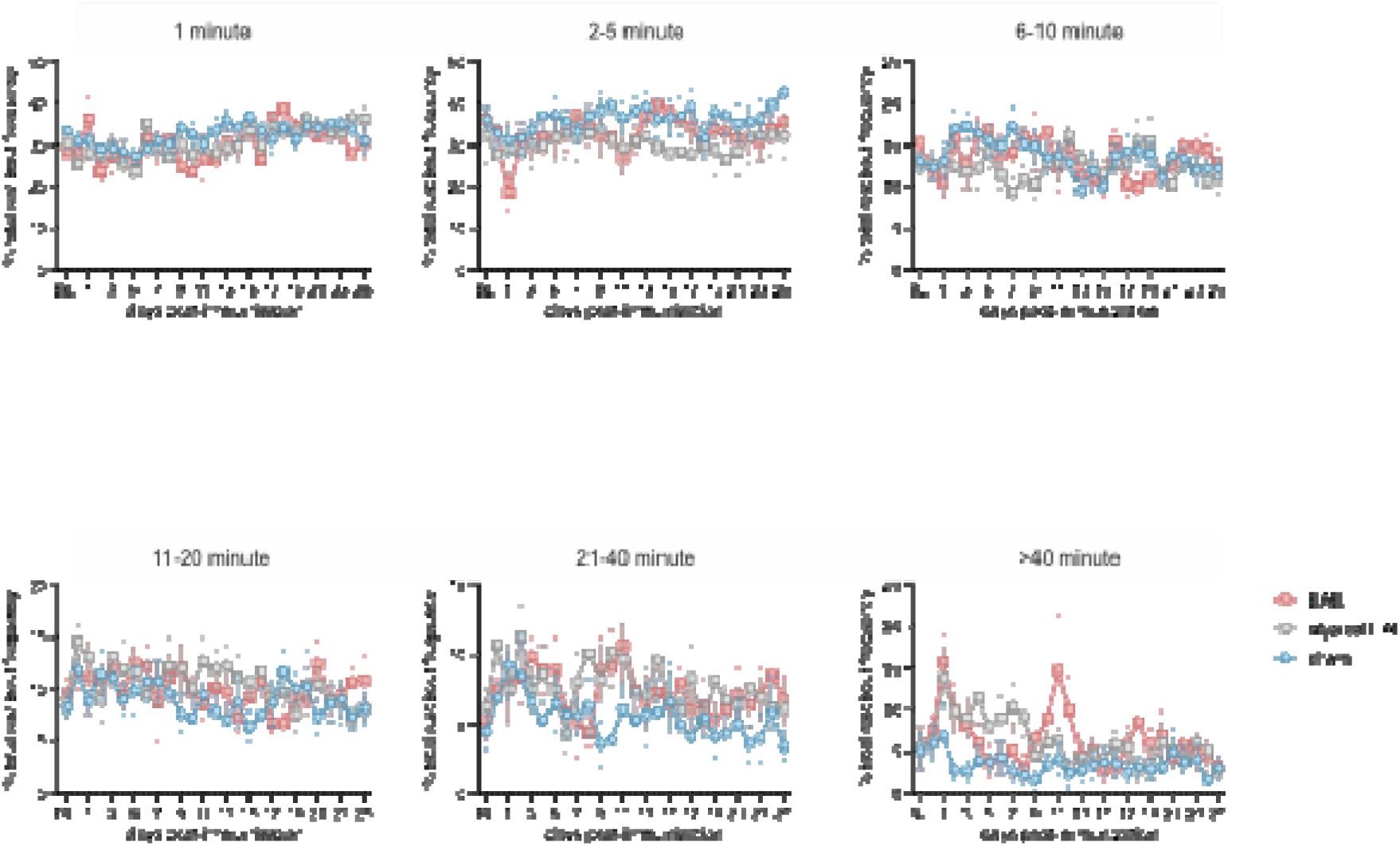
Daily normalized frequency of rest bouts at various lengths in EAE, atypical EAE and sham mice. All data analyzed with two-way RM ANOVA with post-hoc Tukey’s test; n = 5-6 cages/group, 2 animals/cage; ∗p < 0.05, ∗∗p < 0.01, ∗∗∗p < 0.001, ∗∗∗∗p < 0.0001; all data are represented as mean ± SEM. ∗ EAE vs sham, ^#^ atypical EAE vs sham, ^$^ EAE vs atypical EAE.

**Supplementary Figure 10.**
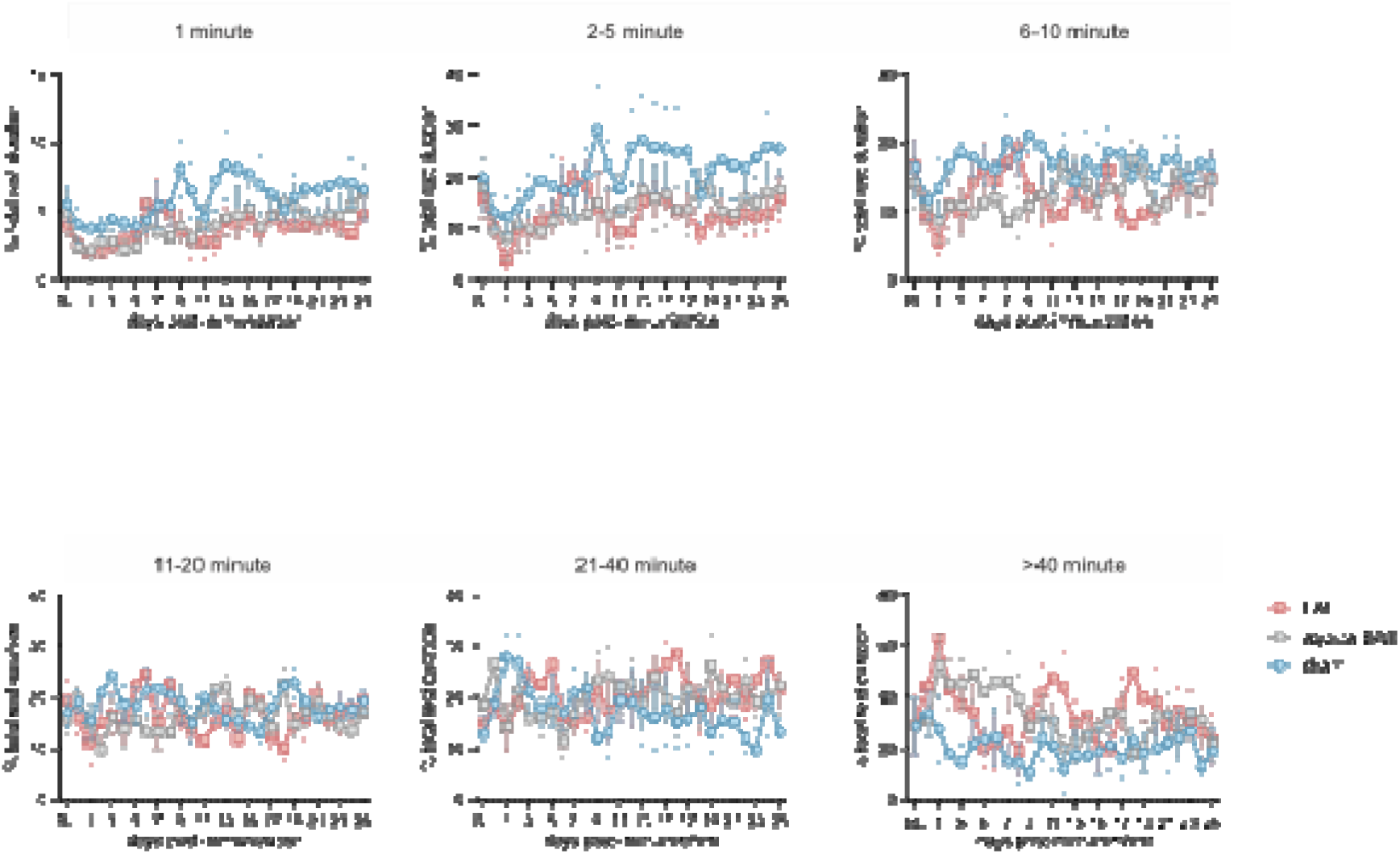
Daily normalized duration of rest bouts at various lengths in EAE, atypical EAE and sham mice. All data analyzed with two-way RM ANOVA with post-hoc Tukey’s test; n = 5-6 cages/group, 2 animals/cage; ∗p < 0.05, ∗∗p < 0.01, ∗∗∗p < 0.001, ∗∗∗∗p < 0.0001; all data are represented as mean ± SEM. ∗ EAE vs sham, ^#^ atypical EAE vs sham, ^$^ EAE vs atypical EAE.

**Supplementary Figure 11.**
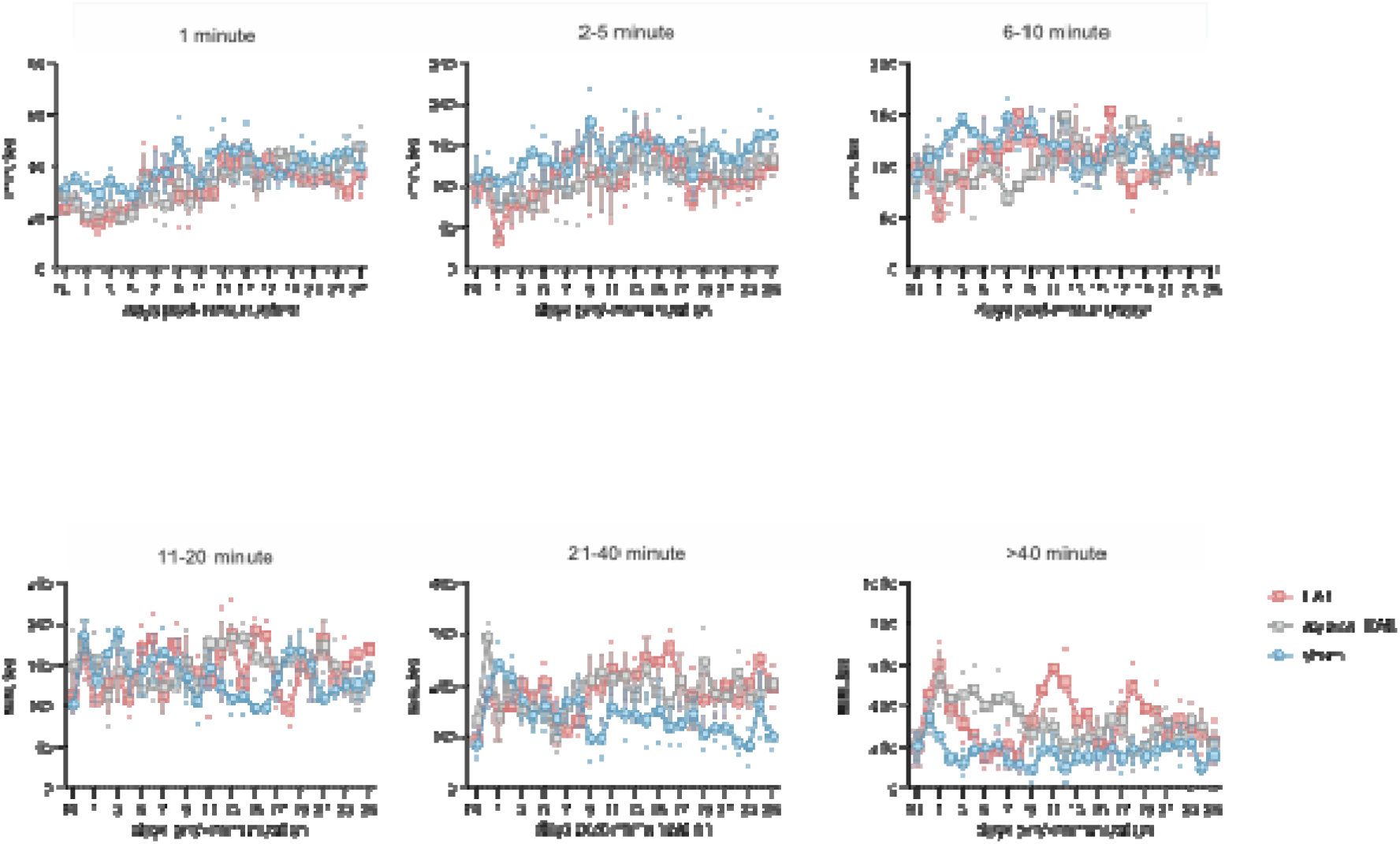
Daily total duration of rest bouts at various lengths in EAE, atypical EAE and sham mice. All data analyzed with two-way RM ANOVA with post-hoc Tukey’s test; n = 5-6 cages/group, 2 animals/cage; ∗p < 0.05, ∗∗p < 0.01, ∗∗∗p < 0.001, ∗∗∗∗p < 0.0001; all data are represented as mean ± SEM. ∗ EAE vs sham, ^#^ atypical EAE vs sham, ^$^ EAE vs atypical EAE.

**Supplementary Figure 12.**
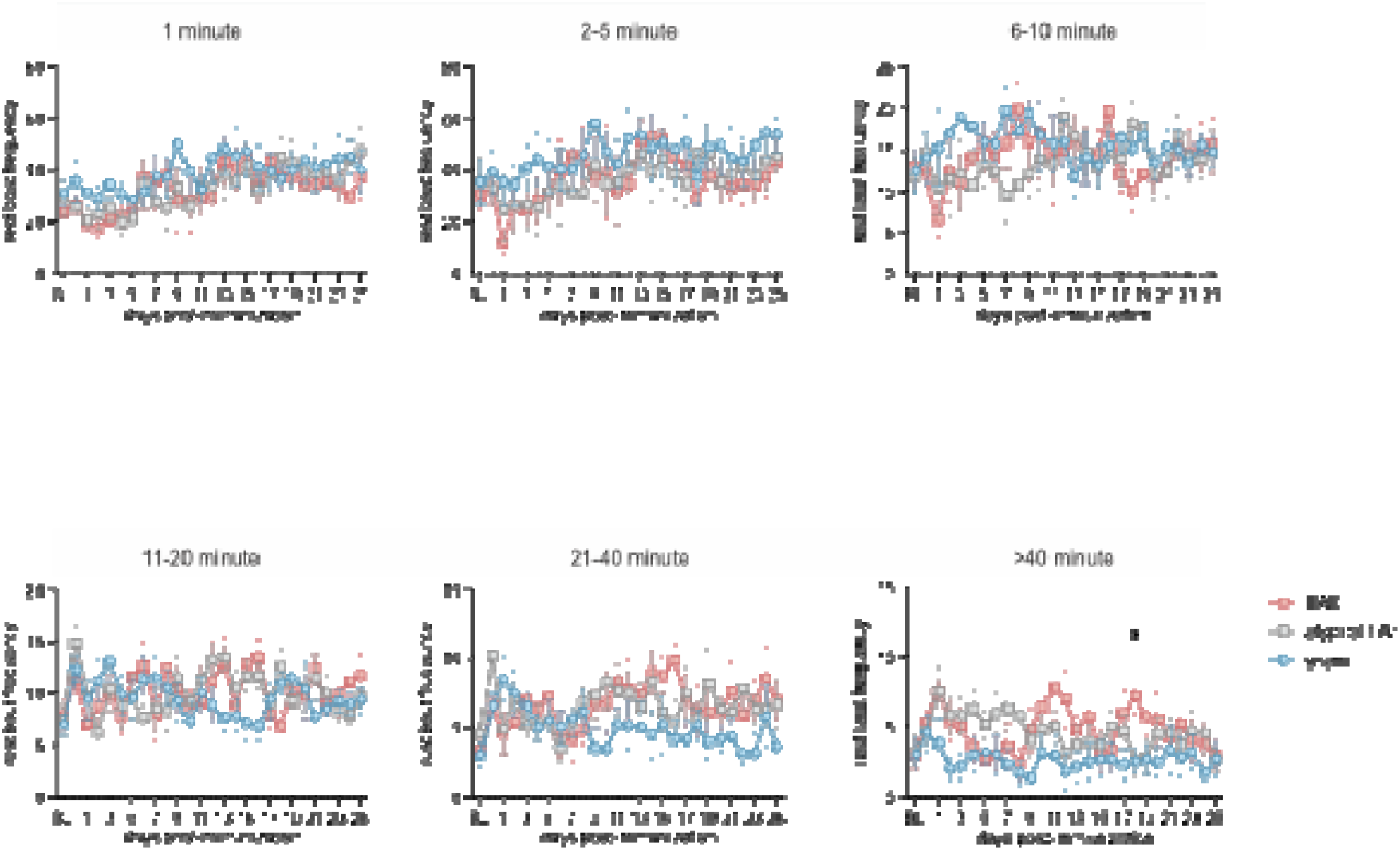
Daily frequency of rest bouts at various lengths in EAE, atypical EAE and sham mice. All data analyzed with two-way RM ANOVA with post-hoc Tukey’s test; n = 5-6 cages/group, 2 animals/cage; ∗p < 0.05, ∗∗p < 0.01, ∗∗∗p < 0.001, ∗∗∗∗p < 0.0001; all data are represented as mean ± SEM. ∗ EAE vs sham, ^#^ atypical EAE vs sham, ^$^ EAE vs atypical EAE.

**Supplementary Table 1.**
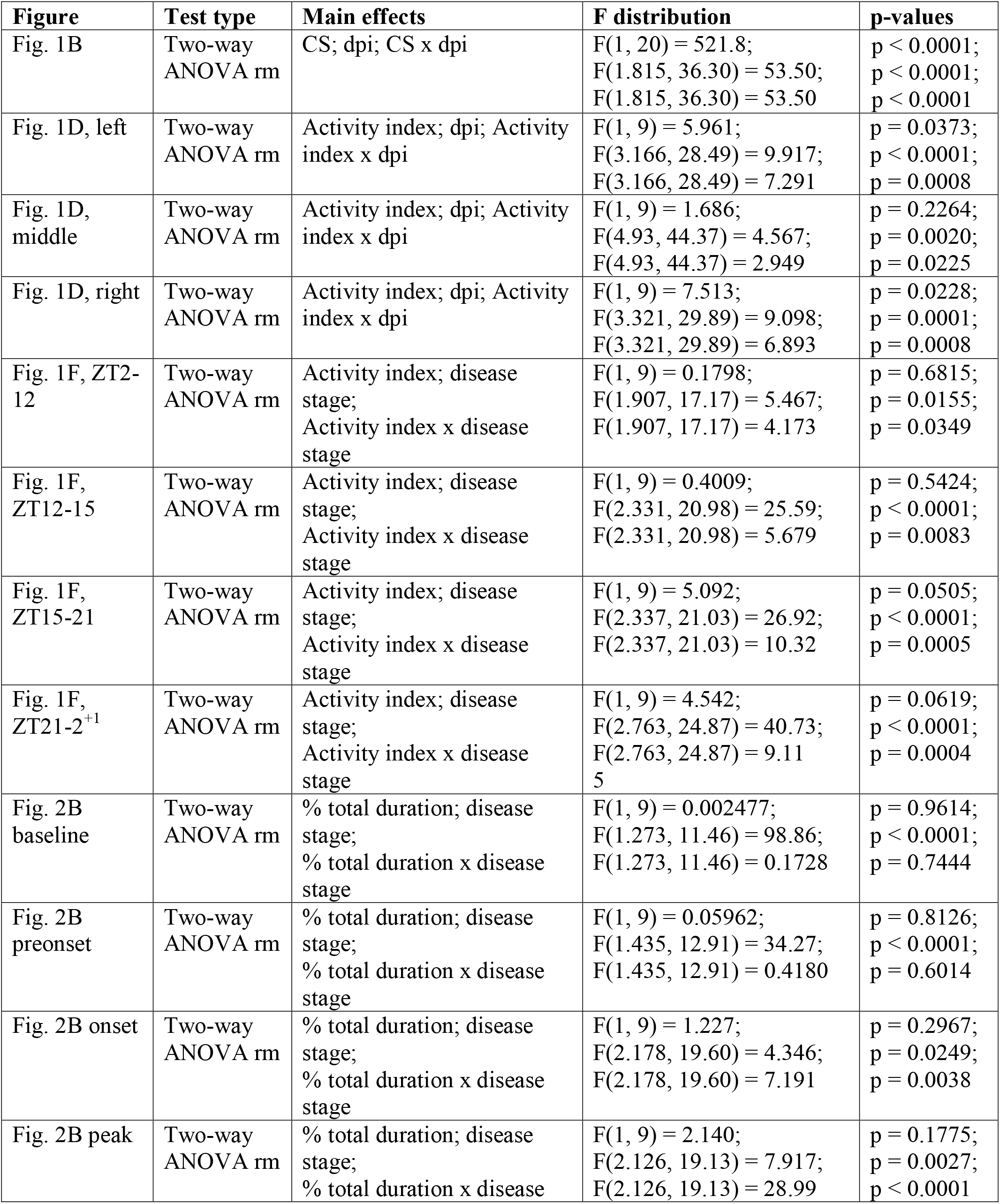
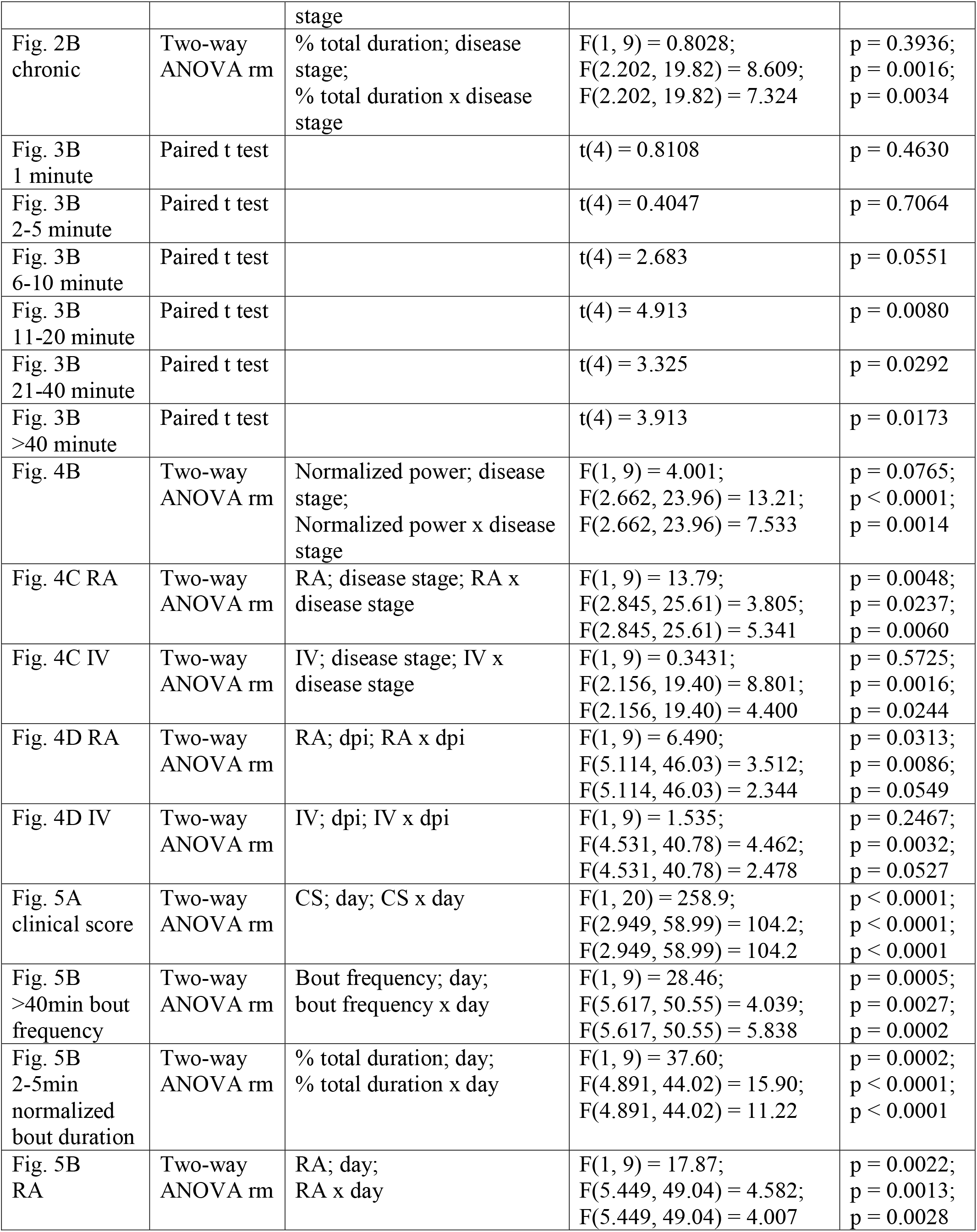
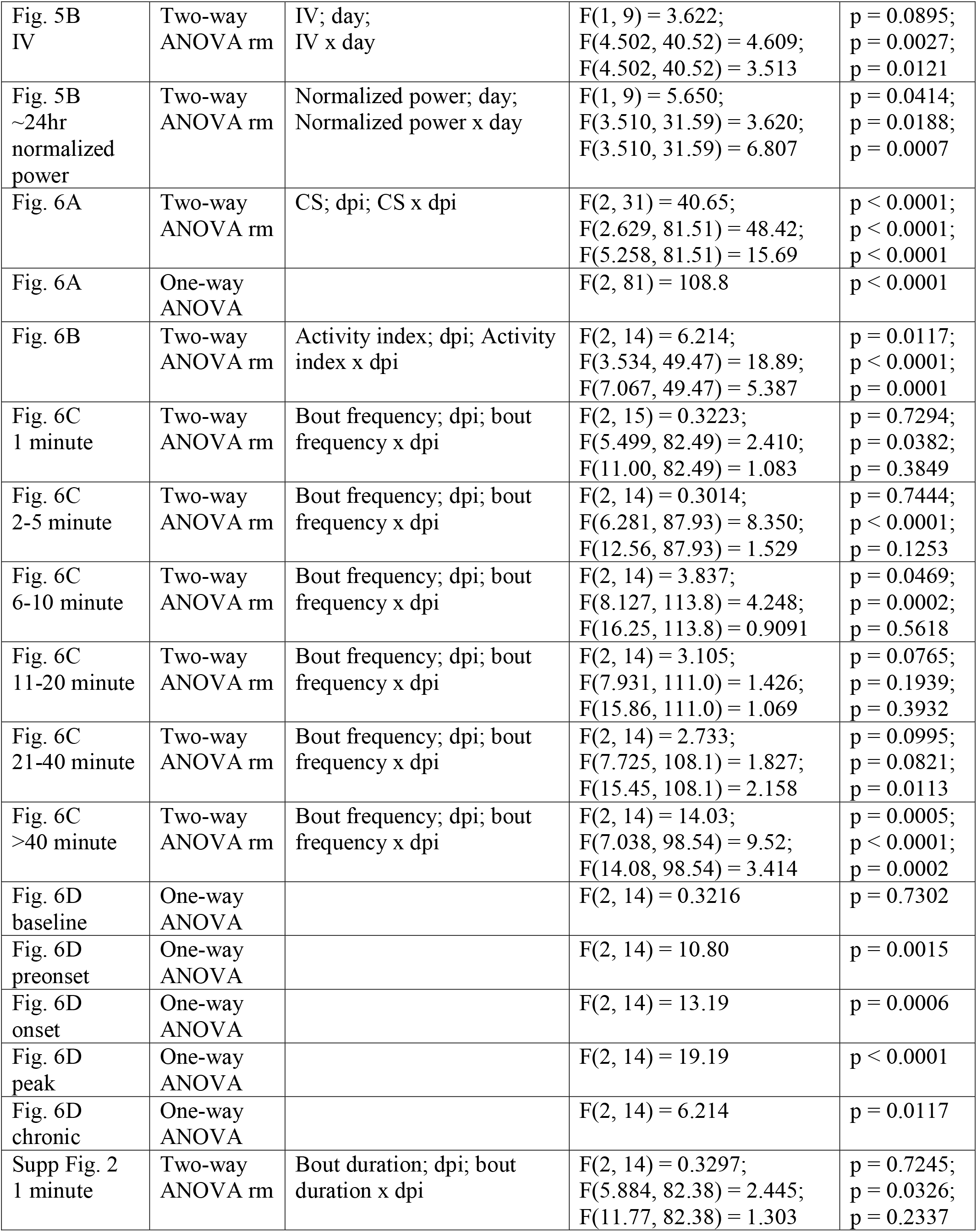
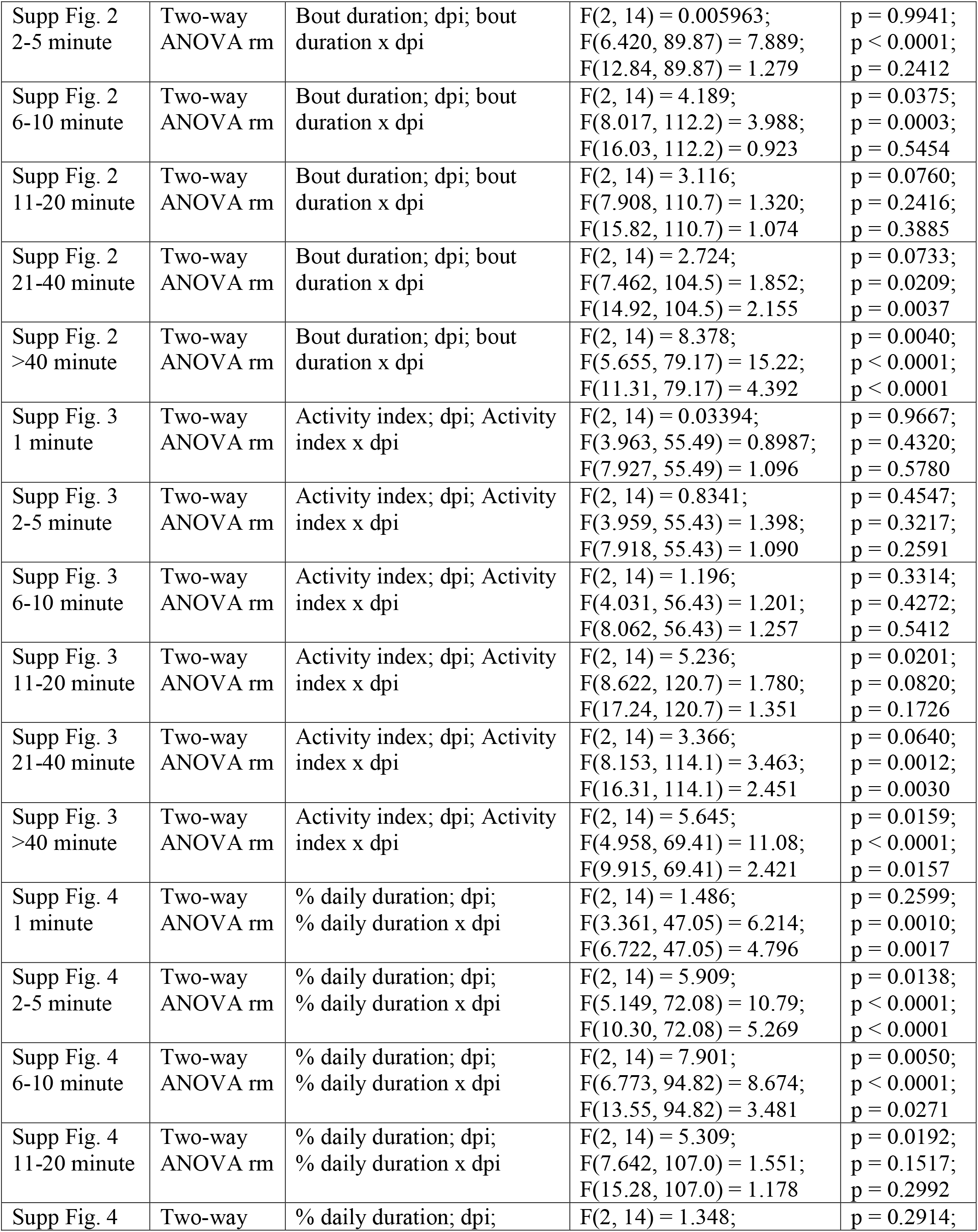
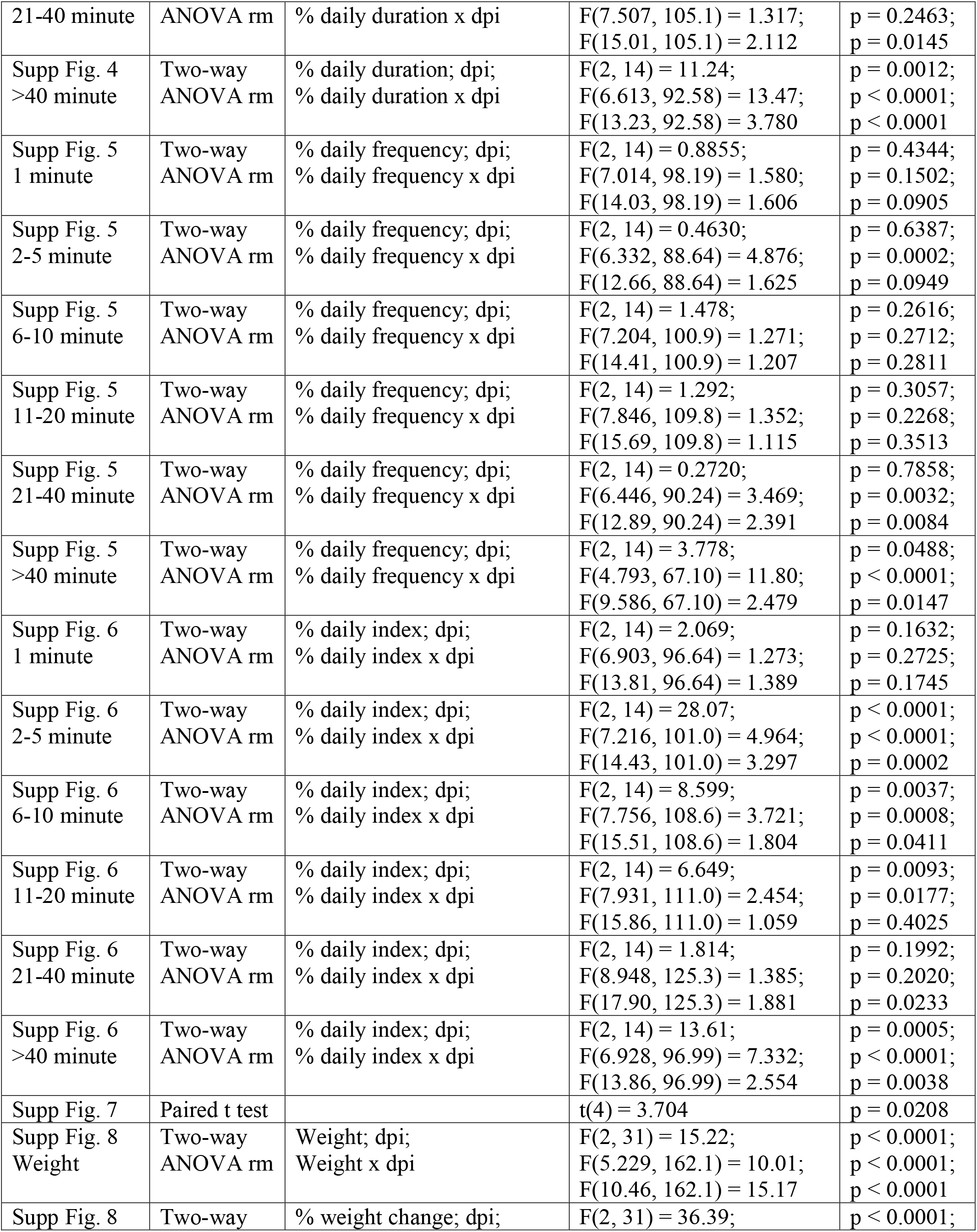
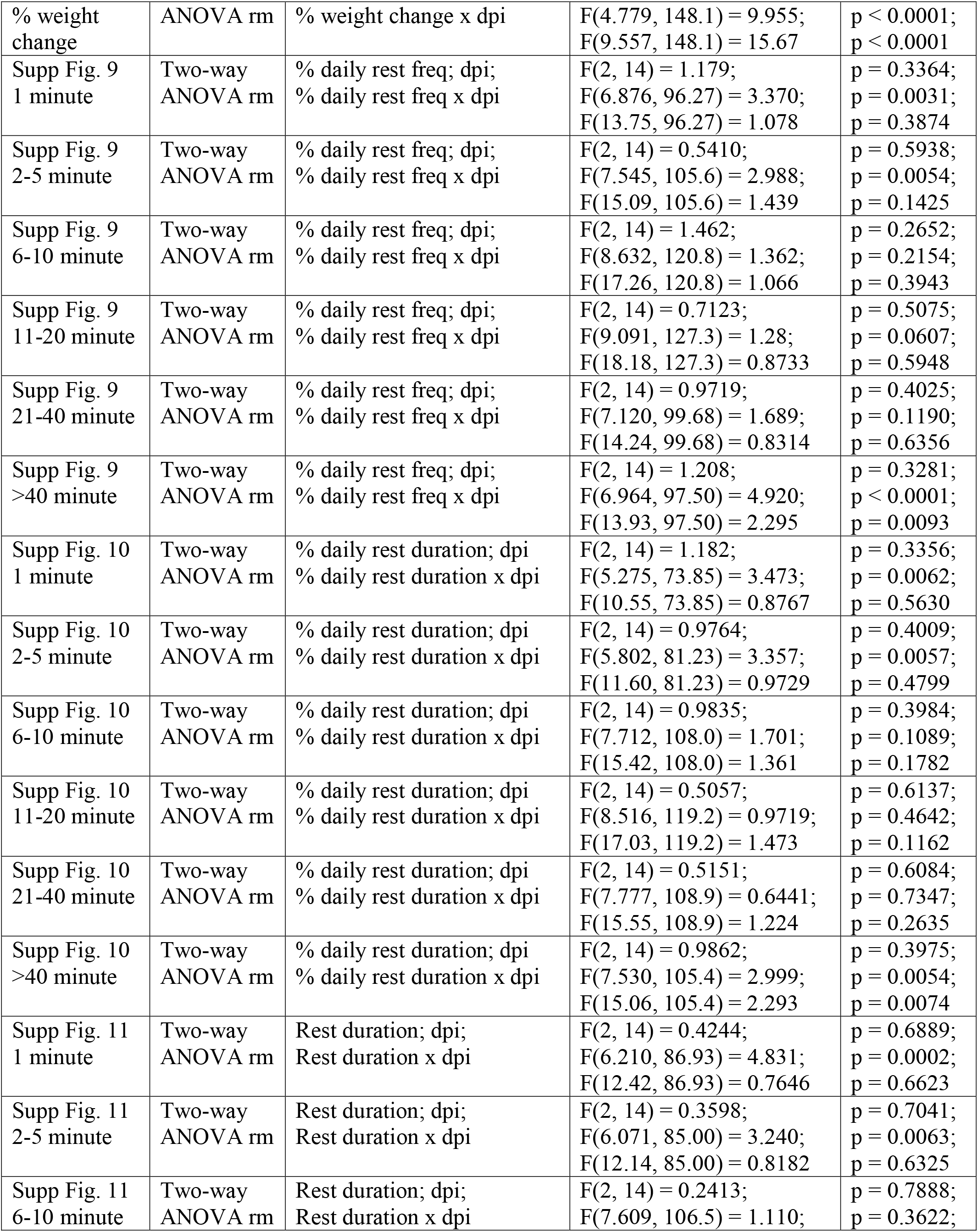
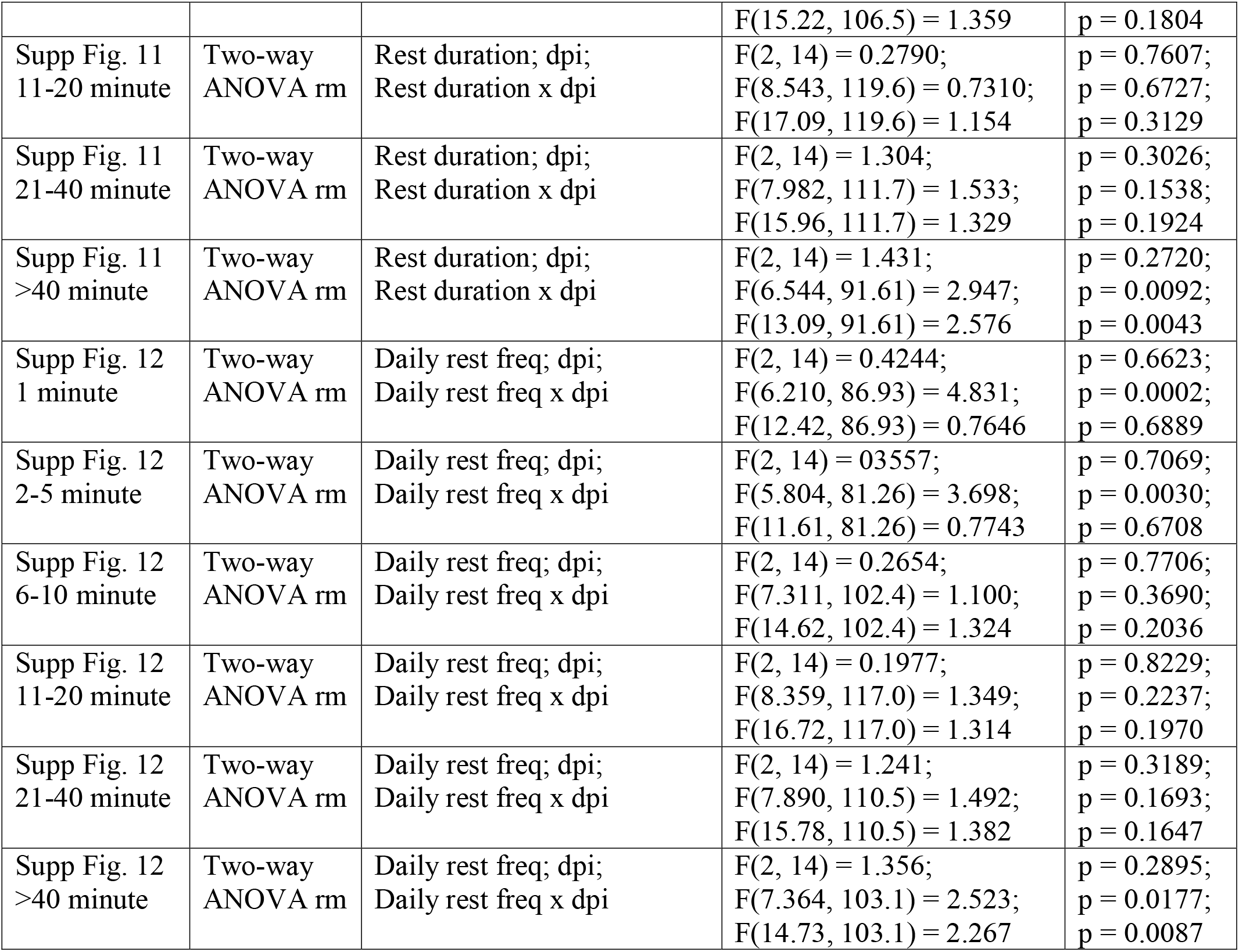
Figure statistical comparison details.

